# Higher-order organization of bacterial cords and an amyloid-like matrix define the *Mycobacterium tuberculosis* biofilm

**DOI:** 10.1101/2025.11.07.687260

**Authors:** Bei Shi Lee, Matthias Godejohann, Richa Mishra, Clément Bousquet, Flavia Gürtler, Abigail J. Deloria, Mengyang Liu, Rainer Leitgeb, Wolfgang Drexler, Gregor L. Weiss, Vivek V. Thacker, Michael Berney, Richard Haindl

## Abstract

Bacterial biofilms generate collective properties that cannot be explained by individual cells alone^1^. Yet the structural principles that organize these communities and their relationship to drug tolerance remain poorly understood^2^, including in major human pathogens such as *Mycobacterium tuberculosis* (*Mtb*)^3,4^. Here we show that submerged *Mtb* biofilms form organized multicellular architectures in which bacterial cords are integrated with a composite extracellular matrix. The virulence lipid phthiocerol dimycocerosate (PDIM) governs the higher-order organization of cords, proteinaceous matrix interactions support cohesion and attachment, an amyloid-like component contributes to biofilm integrity and establishment, and the ESX-1 secretion system increases matrix biochemical complexity. Biofilm growth conferred antibiotic tolerance across genetic backgrounds, while PDIM-dependent organization provided additional protection against first-line anti-tuberculosis drugs, most clearly isoniazid. Targeting amyloid assembly with epigallocatechin gallate impaired biofilm establishment without inhibiting planktonic growth. Together, these findings establish a cording-centred framework linking bacterial surface composition, cellular organization and extracellular matrix assembly to antibiotic tolerance, and show how resolving higher-order biofilm architecture can reveal processes amenable to chemical perturbation.

## INTRODUCTION

*Mycobacterium tuberculosis* (*Mtb*), the causative agent of tuberculosis (TB), is the leading cause of death from a single infectious agent worldwide^5–8^. Its persistence during treatment reflects not only genetic drug resistance but also phenotypic drug tolerance, in which genetically susceptible bacteria survive antibiotic exposure without acquiring heritable resistance^6,9–11^. This tolerant state is frequently associated with altered metabolic activity and environmental stress, and is increasingly linked to growth within multicellular communities^2,3,12–14^. Understanding how such communities are organized, and how their architecture contributes to drug tolerance, is therefore central to understanding bacterial persistence during treatment^1,3,4,12,15–17^.

A hallmark of *Mtb* multicellular behavior is “cording”, the formation of tight, serpentine bacterial bundles linked to virulence^18–21^. Although the glycolipid trehalose dimycolate (TDM) was historically identified as a “cord factor”^22^, the mechanisms underlying cording are more complex^20^. Both the cell-envelope lipid phthiocerol dimycocerosate (PDIM)^20,23,24^ and the ESX-1 secretion system^25–29^ have been implicated in this process. PDIM modulates surface hydrophobicity, whereas ESX-1 mediates the secretion of virulence-associated effectors, suggesting coordinated lipid-protein interactions in shaping multicellular organization.

*Mtb* also forms surface-associated biofilms that have been identified in human lung tissue^4^, yet little is known about their higher-order architecture. Consequently, it remains unclear how individual bacterial assemblies are organized within the biofilm, which molecular components contribute to this organization, and how these features relate to biofilm-associated phenotypes such as drug tolerance.

Here, we investigate how cell-surface determinants and extracellular matrix components organize submerged *Mtb* biofilms using genetically defined strains^30–32^, complementary fluorescence and electron microscopy, label-free quantum cascade laser (QCL)-based mid-infrared microspectroscopy and biochemical perturbation. We distinguish biomass accumulation from higher-order architecture and identify organized multicellular biofilms in which bacterial cords are integrated with a protein-rich extracellular matrix. Our findings support distinct roles for PDIM in the higher-order organization of cords, ESX-1 in matrix biochemical complexity and an amyloid-like component in biofilm integrity and establishment. Functionally, the biofilm state confers antibiotic tolerance across genetic backgrounds, with PDIM-dependent organization providing additional protection against isoniazid. Together, these results establish a cording-centred framework linking bacterial surface composition, multicellular architecture and extracellular matrix assembly to antibiotic tolerance.

## RESULTS

### Thiol reductive stress induces submerged *Mtb* biofilms independently of PDIM and ESX-1

Thiol reductive stress (TRS) rapidly induces *Mtb* biofilm formation in vitro^13^. We first asked whether the virulence-associated determinants PDIM or ESX-1 are required for TRS-induced biomass accumulation. To this end, we examined four H37Rv-derived auxotrophic strains based on mc² 6230 or mc² 6206^33–36^ that differed in their ability to produce PDIM and in the presence or absence of the RD1 locus encoding the ESX-1 secretion system (see Methods).

After exposure to 6 mM dithiothreitol (DTT), all four strains formed submerged biofilms within 72 h (Extended Data Fig. 1a). Despite their genetic differences, crystal violet (CV) staining revealed no significant differences in total biomass among strains (Extended Data Fig. 1b).

Thus, PDIM and ESX-1 are dispensable for TRS-induced bulk biomass accumulation, providing a common baseline for examining the architectural and functional differences described below.

### PDIM promotes the higher-order organization of cords in *Mtb* biofilms

While the total biomass of the biofilms was comparable across strains (Extended Data Fig. 1b), their higher-order organization differed markedly. To resolve these differences, we developed a multicomponent staining protocol to simultaneously visualize α-and β-linked polysaccharides together with the nucleic acids of the resident bacilli (Fig. 1).

**Figure 1:**
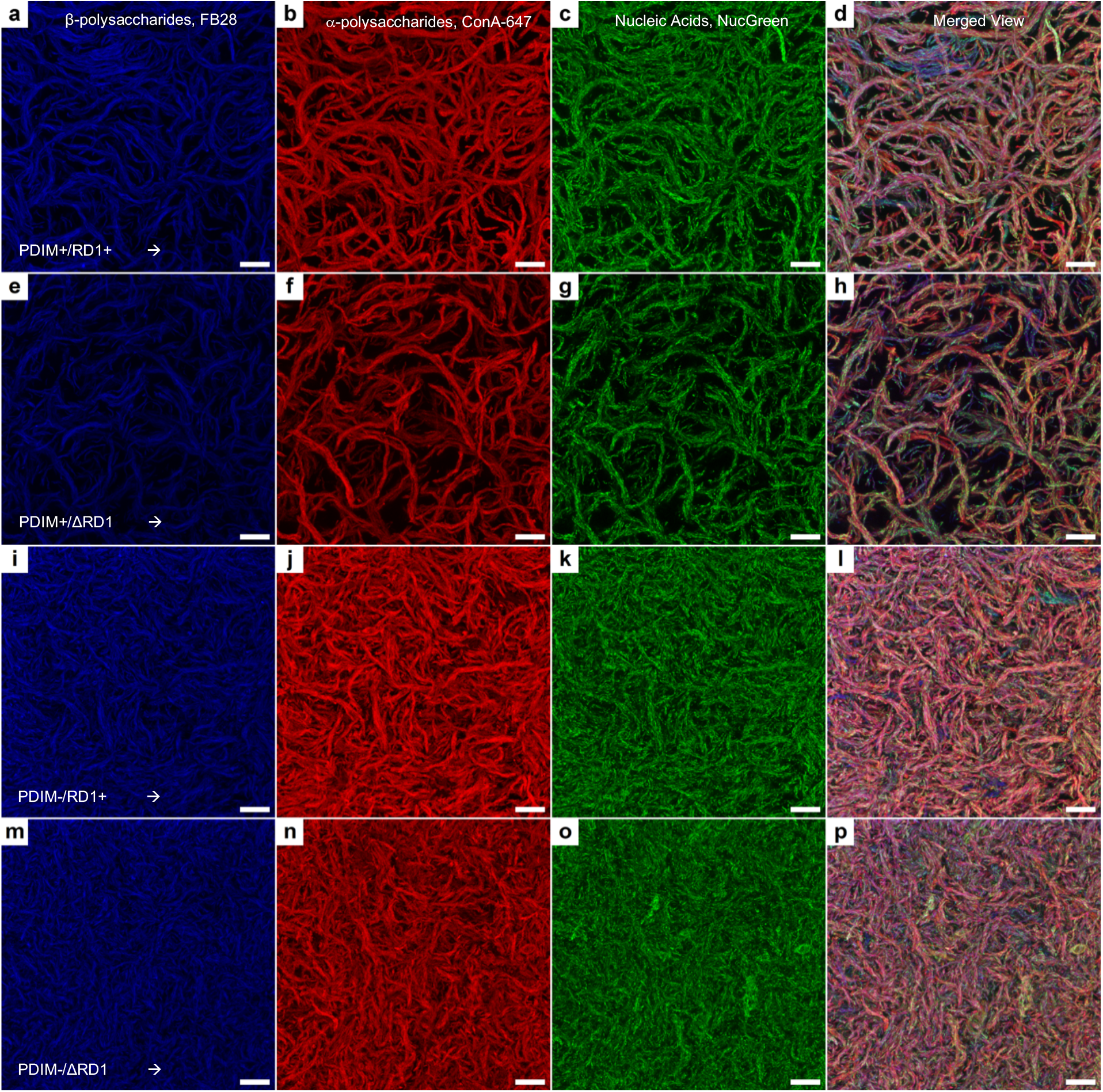
Multicomponent fluorescence imaging reveals genotype-dependent *Mtb* biofilm architectures. Representative confocal microscopy images of paraformaldehyde-fixed biofilms formed by PDIM+/RD1+ (a–d), PDIM+/ΔRD1 (e–h), PDIM−/RD1+ (i– l) and PDIM−/ΔRD1 (m–p) backgrounds. Columns show β-polysaccharides stained with Fluorescent Brightener 28 (FB28, blue), α-polysaccharides stained with concanavalin A Alexa Fluor 647 (ConA-647, red), nucleic acids stained with NucGreen Dead 488 (green) and merged images, respectively. Scale bars, 20 µm.

This approach revealed that all strains produced matrices containing both polysaccharide classes, which co-localized around the bacteria (Fig. 1d, h, l, p). However, the higher-order organization of these matrix components varied substantially between the strains. We selected the α-and β-polysaccharide channels and visualized them as both maximum-intensity projections and 3D volume renderings (Extended Data Fig. 2). In both PDIM-positive backgrounds, PDIM+/RD1+ and PDIM+/ΔRD1, the matrix was organized into elongated, well-defined cord-like structures (Extended Data Fig. 2a, b, e, f). In contrast, PDIM-deficient strains did not form comparably extended, ordered structures. The PDIM−/RD1+ background produced a dense but highly disordered network of shorter, thinner strands (Extended Data Fig. 2c, g), whereas the PDIM−/ΔRD1 background formed a compact, “fluffy” matrix of bacterial aggregates (Extended Data Fig. 2d, h). These data indicate that PDIM is a major determinant of higher-order cord-like organization within the biofilm. The comparison between PDIM-positive backgrounds further suggests that ESX-1 contributes to architectural refinement. The PDIM+/RD1+ biofilm appeared denser and more intricate than the PDIM+/ΔRD1 biofilm, despite both retaining PDIM (Extended Data Fig. 2e, f). Thus, while PDIM is required for the emergence of cord-like organization, ESX-1 appears to enhance scaffold consolidation or complexity.

The alignment of NucGreen staining along the cord-like structures suggested that these features contained organized bacilli. To directly confirm this, we performed transmission electron microscopy on formaldehyde-fixed biofilms. In PDIM-positive backgrounds, bacilli were arranged in tightly aligned bundles within these structures (Extended Data Fig. 3a). Cross-sectional views revealed cords composed of five to six closely packed bacteria across the diameter, with uniform circular profiles consistent with elongated, high-aspect-ratio assemblies. In contrast, bacilli in PDIM-deficient biofilms did not display the extended, tightly aligned bundles observed in PDIM-positive backgrounds and instead appeared more dispersed or in smaller aggregates (Extended Data Fig. 3b). The morphology and organization of these structures are consistent with the classical description of *Mtb* cording in pellicles and intracellular infections^3,20,21^. Together, these data confirm that the cord-like structures observed in biofilms contain aligned bacilli and represent bona fide cords.

### Mid-infrared imaging reveals ESX-1-dependent biochemical complexity within the biofilm matrix

While fluorescence imaging resolved the physical architecture of the biofilms, it provided little insight into their molecular composition. To map the chemical organization of the matrix, we applied quantum cascade laser (QCL)-based mid-infrared (mid-IR) microspectroscopy, enabling label-free visualization of biomolecular components through their inherent vibrational absorption signatures^37,38^. After acquiring hyperspectral data cubes, we generated “Thresholded Specific Wavenumber Abundance Maps” (TSWAMs) to visualize the spatial distribution of key chemical features (Fig. 2; see Methods).

**Figure 2:**
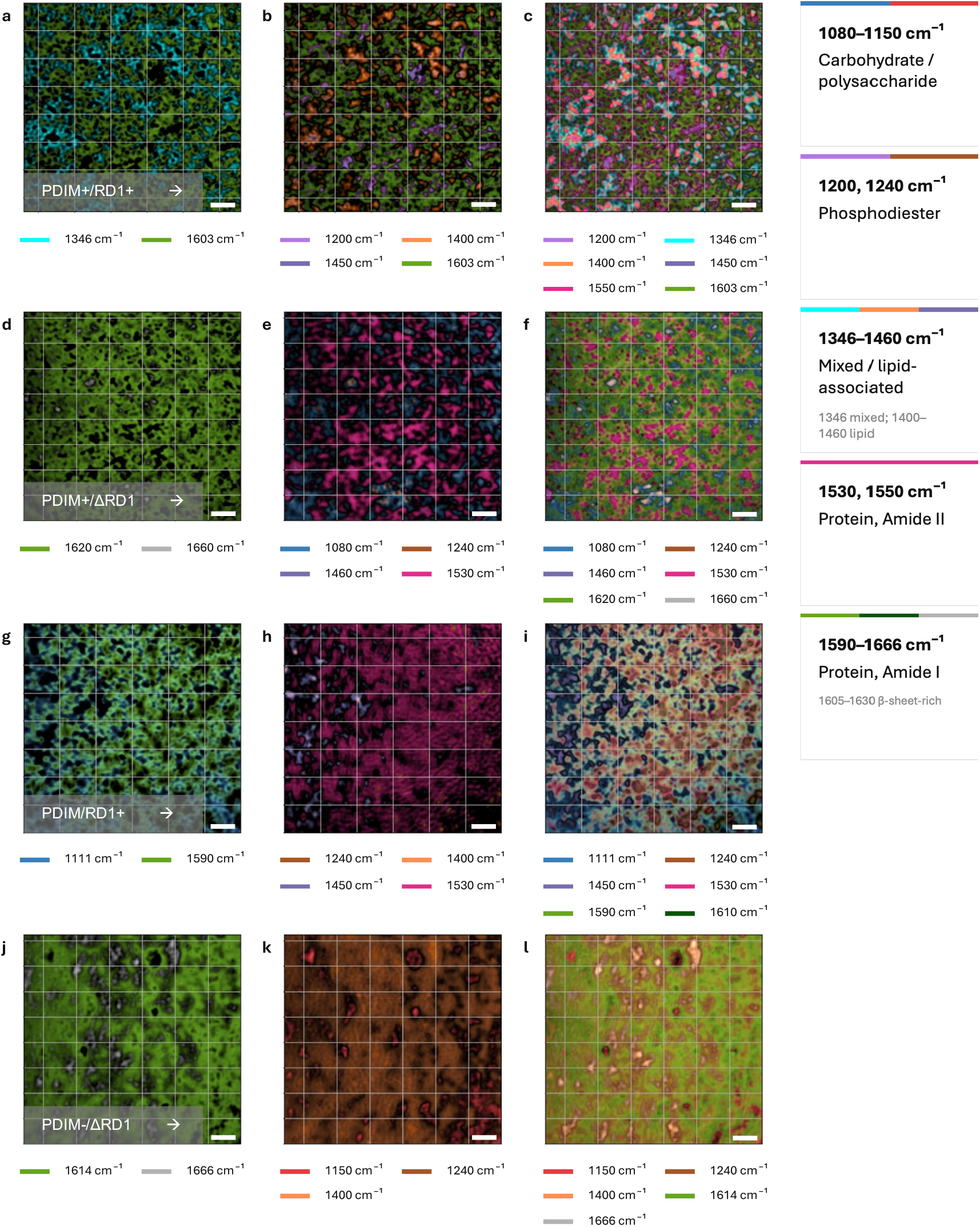
Mid-IR microspectroscopy reveals genotype-dependent biochemical composition and compartmentalization in *Mtb* biofilms. Thresholded Specific Wavenumber Abundance Maps (TSWAMs; Methods) of fixed *Mtb* biofilms imaged in reflection mode. Colours represent the spatial distribution of selected biomolecular signals at the wavenumbers indicated beneath each panel, with broader assignments summarized in the key legend to the right. Rows correspond to PDIM+/RD1+ (a– c), PDIM+/ΔRD1 (d–f), PDIM−/RD1+ (g–i) and PDIM−/ΔRD1 (j–l) biofilms. From left to right, successive addition of selected wavenumber channels provides composite visualizations of the biochemical architecture. Scale bars, 200 µm.

Across all genetic backgrounds, TSWAMs revealed a prominent signal from the protein backbone within the amide I region (1600–1620 cm⁻¹; Fig. 2a, d, g, j), indicating a matrix enriched in proteinaceous material. In the PDIM+/RD1+ biofilm, the principal protein-associated peak occurred at 1603 cm⁻¹, with additional contributions at 1346 cm⁻¹ (Fig. 2a). Successive inclusion of phosphodiester-associated (1200 cm⁻¹) and lipid-associated (1450 and 1400 cm⁻¹) signals progressively filled the spatial map, while an additional protein-associated signal at 1550 cm⁻¹ revealed a highly compartmentalized architecture spanning most of the biofilm (Fig. 2b, c).

This compartmentalization was reduced in the PDIM+/ΔRD1 biofilm. The primary protein-associated signal (∼1620 cm⁻¹) remained evident, but regions that were chemospatially distinct in the PDIM+/RD1+ background were instead largely occupied by polysaccharide (∼1080 cm⁻¹) and protein (∼1530 cm⁻¹) signals, resulting in diminished fine-scale compartmentalization (Fig. 2d–f).

PDIM-deficient strains exhibited further divergence in chemical organization. The PDIM−/RD1+ biofilm displayed a matrix dominated by an amide II signal at 1530 cm⁻¹, with lipid (∼1450 cm⁻¹) and phosphodiester (∼1240 cm⁻¹) contributions. Notably, the spatial distribution of its primary protein signal at 1590 cm⁻¹ overlapped extensively with that of a carbohydrate-associated signal at 1111 cm⁻¹, consistent with a protein-carbohydrate composite scaffold (Fig. 2g–i). The PDIM−/ΔRD1 biofilm showed the most pronounced divergence, retaining a protein backbone (∼1614 cm⁻¹) but exhibiting a simplified chemical map dominated by a ubiquitous phosphodiester signal at 1240 cm⁻¹ and minimal spatial compartmentalization (Fig. 2j–l).

To further resolve these chemospatial patterns, we applied k-means clustering to the hyperspectral datasets (Fig. 3, see Methods). Across all genetic backgrounds, QCL-based mid-IR microspectroscopy revealed prominent antiparallel β-sheet-associated signatures (∼1605–1630 cm⁻¹ and ∼1680 cm⁻¹), together with weaker parallel β-sheet contributions (∼1640 cm⁻¹)^39–41^, particularly in the RD1-intact backgrounds. Such matrix-associated β-sheet enrichment is characteristic of cross-β amyloid assemblies^42–44^, providing an initial label-free indication of amyloid-like material within the *Mtb* biofilm matrix.

**Figure 3:**
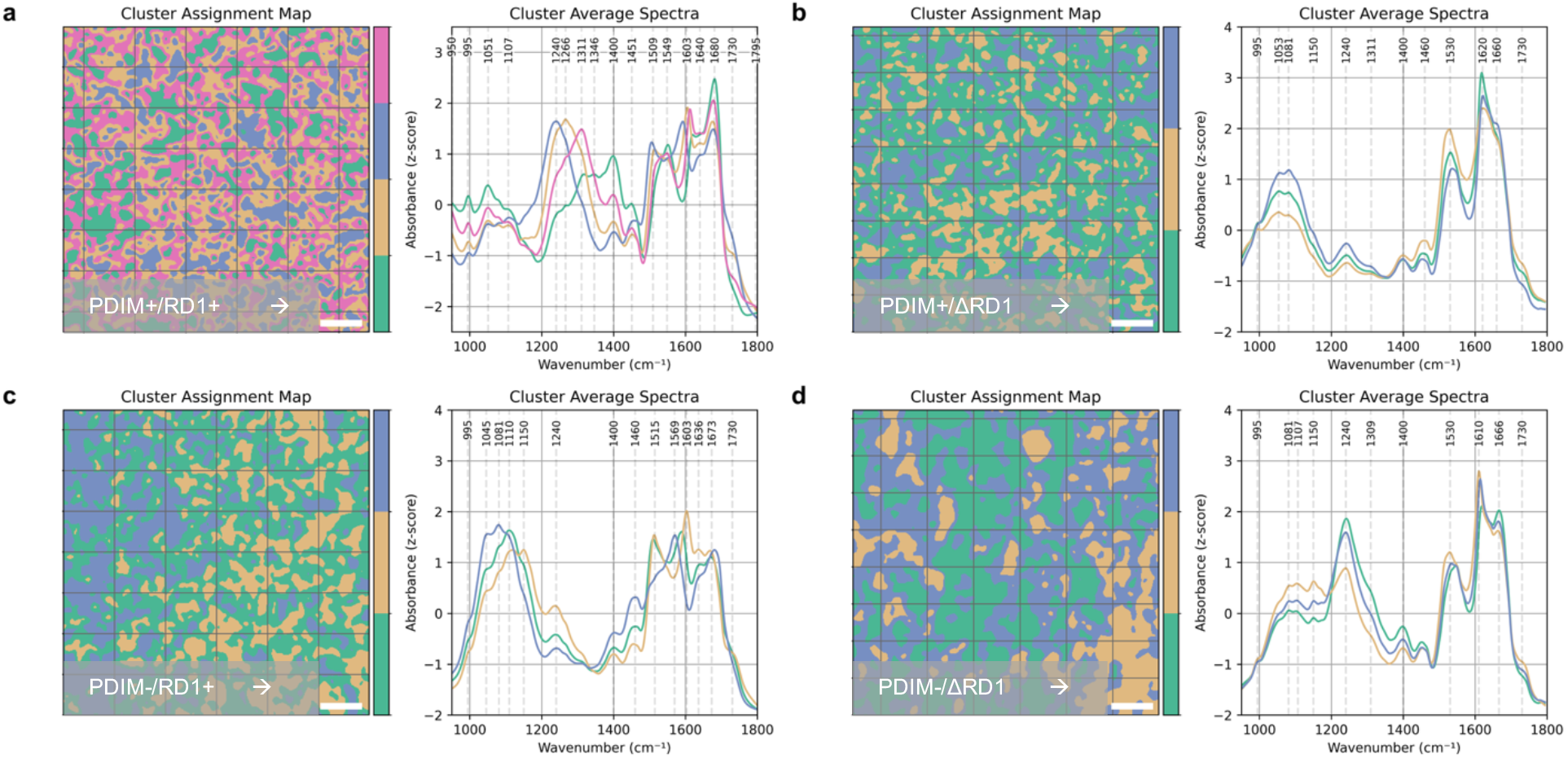
Unsupervised clustering resolves the biochemical architecture of *Mtb* biofilms. **a-d**, K-means segmentation of mid-IR hyperspectral images from PDIM+/RD1+ (a), PDIM+/ΔRD1 (b), PDIM−/RD1+ (c) and PDIM−/ΔRD1 (d) biofilms. For each genetic background, the cluster assignment map (left) shows the spatial distribution of chemospatial regions, with colours denoting distinct clusters. Corresponding cluster-average, z-score-normalized mid-IR spectra are shown on the right using matching colours. Dashed lines and labels identify selected spectral bands. Clustering was performed independently for each background; cluster colours are therefore not directly comparable between panels. Scale bars, 200 µm.

Further, the PDIM+/RD1+ biofilm required four distinct clusters to capture its molecular heterogeneity, with cluster spectra containing diverse protein features and a prominent “mixed signal” region (1250– 1460 cm⁻¹) (Fig. 3a). In contrast, the PDIM+/ΔRD1 biofilm was described by three clusters with largely overlapping spectral profiles that differed mainly in the relative intensity of shared peaks, indicating reduced biochemical diversity (Fig. 3b).

Similarly, the PDIM−/RD1+ biofilm exhibited a simplified chemical profile, with diminished contributions from the mixed signal region. Instead, carbohydrate-associated signals were prominent, with z-scores approaching those of the protein bands (Fig. 3c). The PDIM−/ΔRD1 biofilm showed minimal spatial complexity together with a striking enrichment of phosphodiester-associated signal (∼1240 cm⁻¹) across multiple clusters (Fig. 3d). This phosphodiester signature aligns with the DNA-rich regions observed by fluorescence microscopy (Fig. 1o, p), suggesting that extracellular DNA may assume a more prominent structural role in the absence of both PDIM-dependent organization and ESX-1-driven biochemical maturation. Together, these findings indicate that ESX-1 increases the biochemical complexity of the *Mtb* biofilm matrix.

### An amyloid-like proteinaceous matrix supports biofilm integrity

To identify the key structural components of the biofilm, we subjected biofilms to enzymatic digestions (Fig. 4a, b). Proteinase K treatment caused complete detachment of the biofilms across all four genetic backgrounds, demonstrating that biofilm integrity critically depends on a proteinaceous scaffold, which is consistent with the prominent protein signatures detected by mid-IR microspectroscopy (Fig. 3). Microscopic examination of the detached biofilm material revealed that individual cords were retained, indicating that proteinaceous interactions support biofilm attachment and cohesion between cords rather than bacterial association within individual cords (Extended Data Fig. 4). Negative-stain EM revealed dense extracellular matrix material in native biofilm samples and unbranched fibrils in the supernatant recovered after proteinase K treatment (Fig. 4c). The recovered fibrils had a mean diameter of 4.1 ± 0.5 nm and a mean length of 47.7 ± 14.4 nm (mean ± s.d.; n = 30 fibrils). Their unbranched morphology and dimensions were consistent with those of amyloid fibrils^44^. In contrast, DNase, pullulanase (commercial type I/type II preparation targeting α-1,6 and α-1,4 glucosidic linkages) and cellulase did not measurably reduce retained biofilm biomass (Fig. 4a, b). Nevertheless, pullulanase and cellulase released substantial quantities of reducing sugars, expressed as glucose equivalents, confirming the presence of accessible polysaccharide substrates (Fig. 4d).

**Figure 4:**
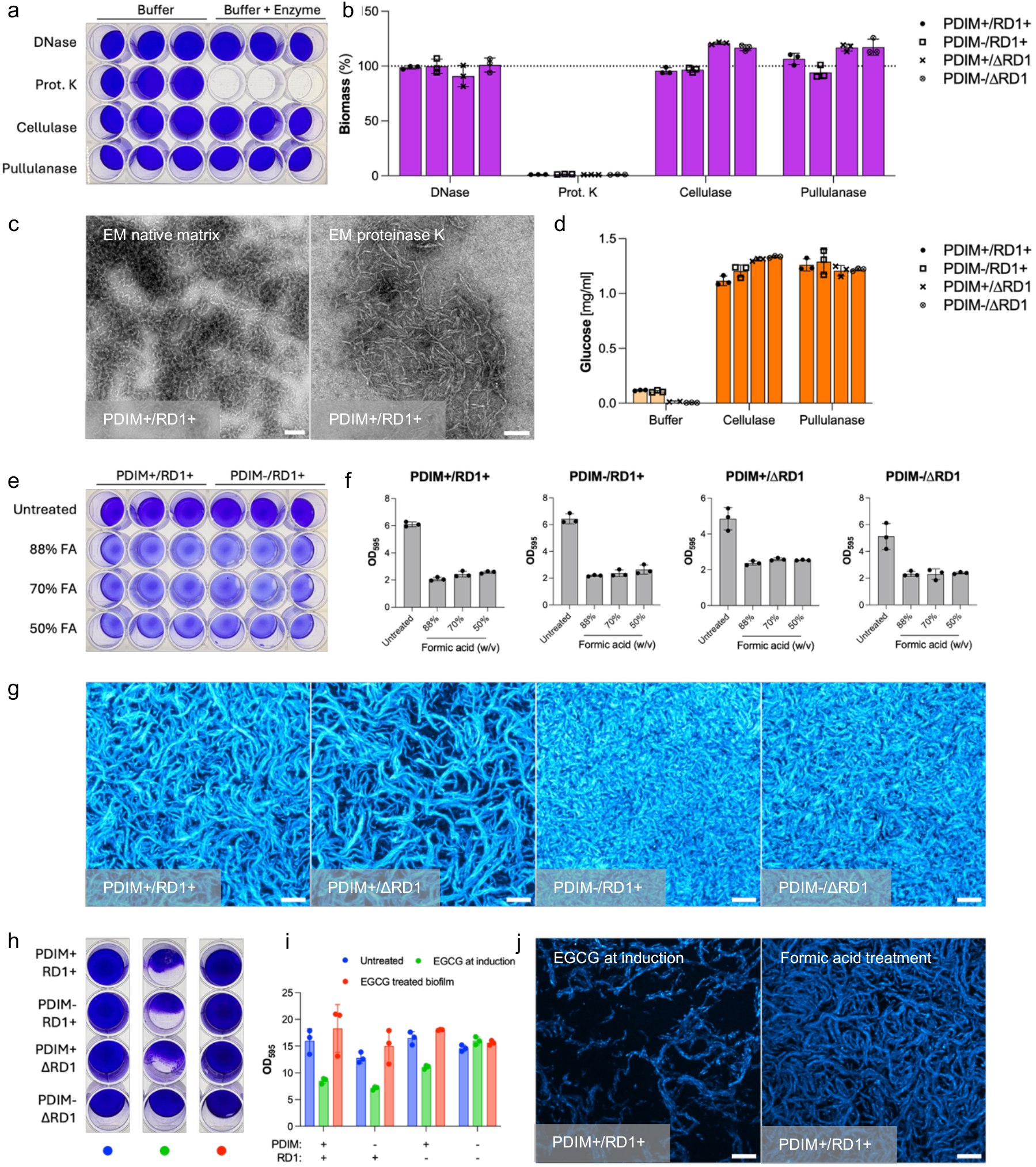
An amyloid-like proteinaceous matrix supports *Mtb* biofilm integrity and formation. a, Representative crystal violet (CV)-stained PDIM+/RD1+ biofilms after enzymatic treatment and corresponding buffer controls. **b,** Biomass after enzymatic treatment across the four genetic backgrounds, normalized to buffer controls. **c,** Negative-stain TEM of native extracellular matrix from PDIM+/RD1+ biofilms (left) and fibrillar material recovered from the supernatant after proteinase K treatment (right). **d,** Reducing sugars released by cellulase or pullulanase, expressed as glucose equivalents. **e,** Representative CV-stained PDIM+/RD1+ and PDIM−/RD1+ biofilms after brief formic acid treatment. **f,** Retained biomass after formic acid treatment across the four genetic backgrounds. **g,** Maximum-intensity projections of ThT fluorescence across the four genetic backgrounds. **h,** Representative CV-stained biofilms left untreated (blue), exposed to EGCG during induction (green) or treated after establishment (red). **i,** Quantification of retained biomass following EGCG treatment either during biofilm induction or after biofilm establishment across the four genetic backgrounds. **j,** Maximum-intensity projections of ThT fluorescence in PDIM+/RD1+ biofilms following EGCG exposure during biofilm induction (left) or formic acid treatment after biofilm establishment (right). Images were acquired using identical confocal settings and displayed over the same intensity range. Depth-resolved analysis is shown in Extended Data Fig. 6. Scale bars, 100 nm (c) and 20 µm (g,j). For b,d,f and i, bars show means, error bars indicate s.d. and symbols represent individual biological replicates (n = 3 per condition).

The proteinase K sensitivity of the biofilm, together with the extracellular fibrils and prominent antiparallel β-sheet-associated spectral signatures detected by mid-IR analysis, prompted us to investigate whether amyloid-like structures contribute to biofilm stability. Brief formic acid treatment, which can disaggregate amyloid assemblies^45^, substantially reduced retained biofilm biomass across all genetic backgrounds and concentrations tested (Fig. 4e, f). To examine the distribution of amyloid-like material, biofilms were stained with the amyloid-binding dye Thioflavin T (ThT)^46^. Confocal microscopy revealed strong ThT-positive material associated with bacterial cords or aggregates across all genetic backgrounds (Fig. 4g).

Together, the fibrillar morphology, β-sheet-associated spectral signatures, formic acid sensitivity and ThT-positive staining support the presence of a proteinaceous scaffold with amyloid-like properties that contributes to *Mtb* biofilm integrity.

Finally, to determine whether these architectural features extend beyond the H37Rv-derived auxotrophic strain panel, we examined *M. tuberculosis* Erdman under the same reductive stress conditions. Wild-type Erdman formed biofilms containing organized cord-like structures and ThT-positive matrix material, whereas the PDIM-deficient Erdman 5’Tn::fadD26 mutant^47,48^ formed disordered aggregates while retaining ThT-positive matrix material (Extended Data Fig. 5). Thus, cord-like organization and ThT-positive matrix material were also observed in a virulent *Mtb* background and were not confined to the auxotrophic strains.

### EGCG impairs *Mtb* biofilm formation

Given the convergent evidence for an amyloid-like matrix component, we next examined whether interference with amyloid assembly affects biofilm formation. To test this, we used (-)-epigallocatechin gallate (EGCG), a polyphenol compound with established anti-amyloidogenic activity^49^. EGCG exhibited no detectable inhibitory effect on planktonic *Mtb* growth in vitro at concentrations up to 250 µg ml⁻¹ (Extended Data Fig. 7), allowing us to assess effects on biofilm formation in the absence of detectable direct antibacterial activity. EGCG was therefore added either at the time of biofilm induction together with DTT or after biofilm maturation following 72 h of reductive stress exposure. Biofilm biomass was quantified four days later with crystal violet staining (Fig. 4h, i). Addition of EGCG during biofilm induction impaired biofilm formation in all genetic backgrounds except PDIM−/ΔRD1. In contrast, treatment of pre-established biofilms produced no detectable reduction in crystal violet staining over the experimental time course.

Although biofilms formed in the presence of EGCG retained measurable crystal violet-positive biomass, maximum-intensity projections revealed a pronounced reduction in ThT-positive material (Fig. 4j). Across the analysed volume, EGCG reduced the ThT-positive volume fraction to 35% and the volume-normalized ThT signal to 13% of untreated levels, with significant differences across the complete depth profiles (Extended Data Fig. 6). Formic acid treatment produced a distinct response: ThT-positive volume fraction remained at 95% of untreated levels, whereas the volume-normalized signal decreased to 45%. The remaining fluorescence was preferentially retained near the glass substratum and decreased towards the exposed biofilm surface. Together, the reduction in ThT fluorescence following both treatments further supports the presence of an amyloid-containing matrix component and indicates that EGCG-mediated impairment of biofilm formation is accompanied by reduced assembly of ThT-positive matrix material.

### Biofilm architecture contributes to antibiotic tolerance

To determine whether differences in biofilm architecture influence antibiotic protection, biofilms were exposed to high concentrations of the frontline antibiotics isoniazid (INH) and rifampicin (RIF), and viability was monitored over 9 days (Fig. 5, Extended Data Fig. 8). In parallel, the same genetic backgrounds were treated under planktonic growth conditions in the absence of DTT and in the presence of tyloxapol to distinguish biofilm-specific effects from intrinsic drug susceptibility (Extended Data Fig. 8).

**Figure 5:**
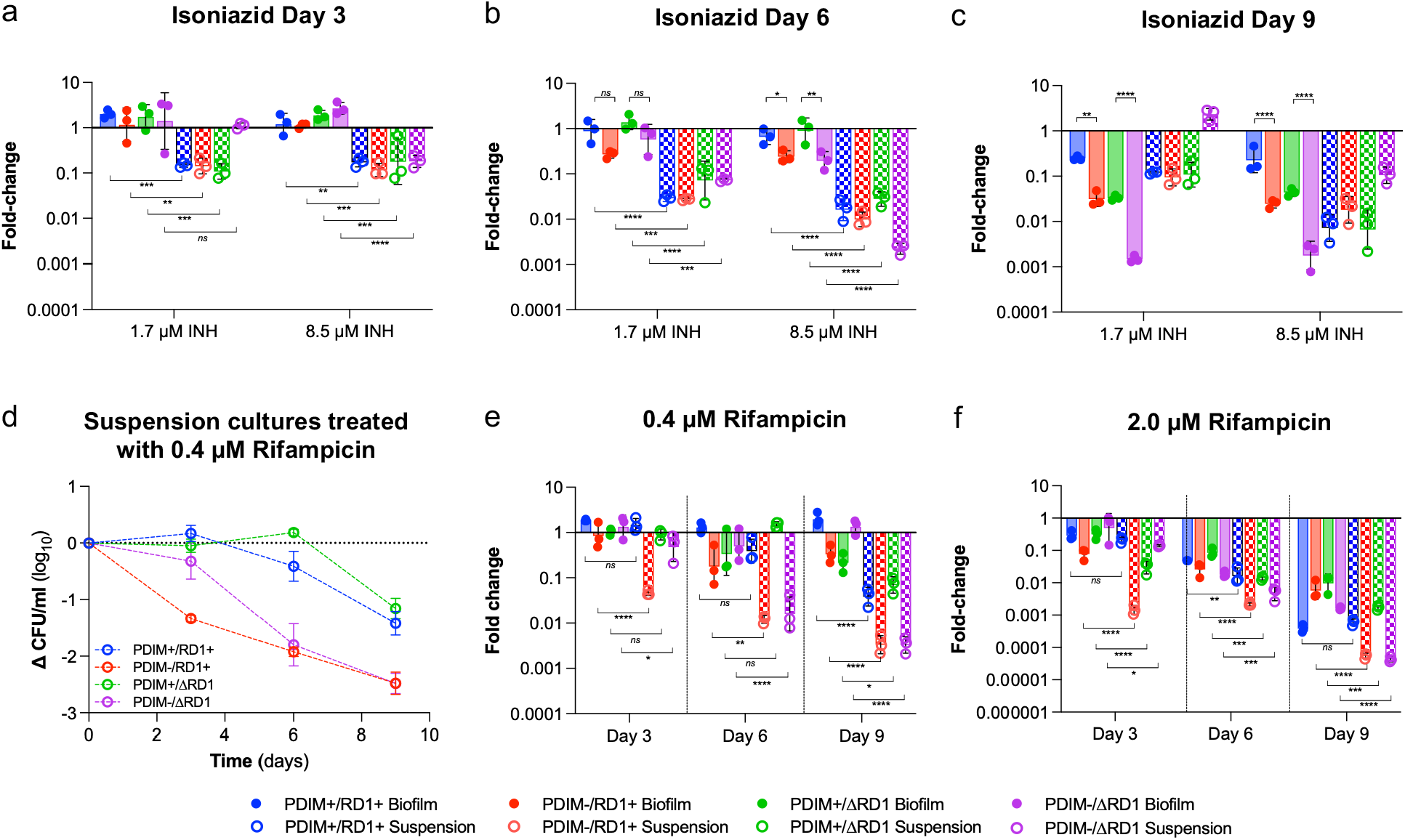
Biofilm architecture contributes to antibiotic tolerance in *Mtb*. **a-c,** Fold change in viable bacteria following treatment with 1.7 or 8.5 µM isoniazid (INH) for 3 days (a), 6 days (b) or 9 days (c). **d,** Change in suspension-culture CFU relative to day 0 during treatment with 0.4 µM rifampicin (RIF). **e,f,** Fold change in viable bacteria following treatment with 0.4 µM (e) or 2.0 µM RIF (f) for the indicated durations. In a-c and e,f, values are expressed relative to the corresponding untreated control for each genetic background, time point and growth state. PDIM+/RD1+, blue; PDIM−/RD1+, red; PDIM+/ΔRD1, green; and PDIM−/ΔRD1, purple. Filled bars and symbols denote biofilm cultures; hatched bars and open symbols denote suspension cultures. Bars show means and overlaid symbols represent individual biological replicate values (n=3 per condition). Error bars, where shown, denote s.d. Statistical significance in a-c and e,f was assessed using two-way ANOVA with Šídák’s multiple-comparisons test for comparisons between biofilm and suspension conditions, and two-way ANOVA with Tukey’s multiple-comparisons test for comparisons among biofilm strains. Adjusted P values are indicated as ns, P ≥ 0.05; *P < 0.05; **P < 0.01; ***P < 0.001; ****P < 0.0001. Test statistics with confidence intervals, effect sizes, degrees of freedom and exact P values are provided in the Supplementary Note.

INH was tested at 1.7 µM and 8.5 µM, corresponding to approximately 10-fold and 50-fold the MIC₅₀ values of the strains (Extended Data Fig. 9a). As expected, PDIM and RD1 status did not alter intrinsic susceptibility to INH. Under planktonic conditions, all strains exhibited the characteristic rapid decline in viability followed by outgrowth after prolonged exposure consistent with the emergence of escape mutants (Extended Data Fig. 8a-d)^50^. In contrast, biofilm-associated bacteria showed markedly delayed susceptibility to INH (Fig. 5a and b, Extended Data Fig. 8a-d). This protective effect was strongly influenced by PDIM production: biofilms formed by PDIM-positive backgrounds showed substantially greater tolerance to INH (Fig. 5b, c).

Biofilm-associated protection was also observed during RIF treatment (Extended Data Fig. 8e–h). However, interpretation of these effects was complicated by known differences in intrinsic RIF susceptibility associated with PDIM status^51^ (Extended Data Fig. 9b). At fixed concentrations of 0.4 µM (Fig. 5d) and 2.0 µM RIF, PDIM-positive planktonic cultures already displayed increased survival relative to PDIM-deficient strains, reducing the apparent contribution of the biofilm state itself. Nevertheless, biofilm-associated protection became more evident at later time points and was most apparent in the PDIM−/RD1+ and PDIM−/ΔRD1 backgrounds (Fig. 5e, f; Extended Data Fig. 8f, h), where the contribution of intrinsic PDIM-associated tolerance was absent. Together, these findings demonstrate that the *Mtb* biofilm state confers substantial antibiotic tolerance across all genetic backgrounds, with PDIM-dependent higher-order organization further amplifying protection against isoniazid.

## DISCUSSION

The ability of *Mycobacterium tuberculosis* to form resilient biofilms represents a major obstacle in tuberculosis treatment, yet the architectural principles underlying this multicellular state remain poorly defined^4,12,13^. Our data support a cording-centred model of *Mtb* biofilm formation. Rather than accumulating as an amorphous extracellular mass, *Mtb* organizes cords into a PDIM-dependent higher-order superstructure. Proteinaceous matrix interactions support cohesion and attachment, amyloid-like material contributes to biofilm integrity and establishment, ESX-1 increases matrix biochemical complexity, and extracellular glycans are incorporated into the resulting composite scaffold. This reframes *Mtb* biofilm formation from a biomass-accumulation process into an architecture-building process and links PDIM-dependent higher-order organization to additional protection against isoniazid within a broader drug-tolerant biofilm state.

The separation between biomass and architecture provides the strongest support for the model. PDIM-deficient strains formed biofilms of comparable biomass but lacked the organized superstructure of cords observed in PDIM-positive strains, despite retaining TDM production. TDM alone is therefore insufficient for higher-order biofilm architecture, whereas PDIM promotes the organization of cords at the community scale. This extends the classical concept of cording from a microscopic virulence-associated morphology to a structural principle of biofilm organization^21,22,52^.

The maintenance of the superstructure depends on proteinaceous matrix interactions. Among the matrix components tested, only protein digestion disrupted overall biofilm integrity, while individual cords were retained. Proteinaceous interactions are therefore not required to maintain bacterial association within cords but they support biofilm attachment and cohesion between cords. The protein-dependent architecture may include amyloid-like material together with other proteinaceous components acting as tethers or attachment factors.

Convergent evidence from orthogonal approaches supports the presence of amyloid-like matrix materials: mid-IR spectroscopy revealed β-sheet-associated matrix signatures consistent with cross-β amyloid assemblies^39–44^; ThT-positive material was associated with bacterial cords or aggregates^46,53^; and brief formic acid treatment reduced both retained biofilm biomass and ThT fluorescence, consistent with the known formic acid sensitivity of amyloid assemblies^45,54,55^. Functional amyloids are well-established structural components of bacterial biofilms, including CsgA in *Escherichia coli*, FapC in *Pseudomonas*, and TasA in *Bacillus subtilis*^42–44^. The resemblance of the *Mtb* fibrils to fibrils from these characterized systems provides additional evidence for a scaffold with amyloid-like properties within the *Mtb* biofilm matrix. We have not yet identified the protein or proteins that contribute to this amyloid-like material. The *Mtb* protein MTP (Rv3312A), which forms curli-like pili, is one candidate^56^; stress-response or surface-associated proteins with β-sheet-forming potential, including HspX and GroEL, may also contribute^3,14,57^.

The EGCG experiments further supported a role for amyloid-like material during biofilm establishment: EGCG impaired biofilm formation and markedly reduced both the extent and fluorescence intensity of the ThT-positive matrix when present during induction, consistent with the established ability of EGCG to interfere with β-sheet-rich amyloid assembly^49,58^. Together with the formic acid experiments, these findings support a contribution of amyloid-like matrix material to both mature biofilm integrity and biofilm establishment. The PDIM−/ΔRD1 background, which showed reduced sensitivity to EGCG, exhibited a prominent phosphodiester-rich matrix signature. This suggests that *Mtb* can establish biofilms through partially distinct architectural routes in the absence of both PDIM-dependent higher-order organization and ESX-1-dependent matrix maturation.

The ESX-1-dependent matrix maturation is reflected in the greater chemospatial heterogeneity of RD1-intact strains relative to their ΔRD1 counterparts. The biochemical pattern is consistent with transcriptomic data showing ESX-1 upregulation in biofilm-dwelling *Mtb*^13^ and raises the possibility that self-assembling ESX-1 substrates such as EspB contribute to matrix complexity^59^.

Extracellular glycans are incorporated into the matrix as well. Glycosidase treatment released reducing sugars, but without measurably decreasing retained biofilm biomass. This argues against a primarily polysaccharide-driven scaffold and instead supports a composite architecture in which accessible glycans are interwoven with the superstructure of cords and proteinaceous matrix components.

The functional consequence of the described organized biofilm architecture is most evident in drug tolerance. All four genetic backgrounds showed enhanced antibiotic tolerance in the biofilm state. Isoniazid provided the clearest link between architecture and protection because planktonic susceptibility was unaffected by PDIM or ESX-1 status, whereas PDIM-positive biofilms amplified a broader protective state conferred by the biofilm matrix. Consistent with general models of biofilm recalcitrance, this protection is likely multifactorial, involving matrix-dependent restriction or sequestration of antibiotics together with biofilm-associated physiological changes^17,60–63^.

The comparable cord-like morphology and ThT-positive matrix observed in the virulent Erdman strain indicate that these features are not confined to the auxotrophic backgrounds. Extending this analysis to diverse clinical isolates and infected host tissue will reveal how the architectural principles identified here operate across bacterial backgrounds and within the more complex network of host-pathogen interactions that may further shape biofilm organization and drug tolerance^4,15,21,52^.

Beyond the biological model, this work highlights the value of chemospatial imaging for studying bacterial biofilms. Fluorescence microscopy resolved cords, polysaccharides and amyloid-associated staining, whereas QCL-based mid-IR microspectroscopy linked this physical architecture to biomolecular composition without labelling^37,38,64–68^. In particular, mid-IR imaging revealed β-sheet-associated matrix signatures that could not have been inferred from morphology alone and guided subsequent orthogonal testing for amyloid-like material^39–44^. This demonstrates how mid-IR microspectroscopy can function as a discovery tool for biofilm matrix biology, while also emphasizing the need for complementary molecular approaches to define the precise identities of the matrix components^65,69^.

Taken together, our data distinguish biomass accumulation from higher-order architecture and support a cording-centred model linking cell-surface composition and matrix maturation to multicellular organization and antibiotic tolerance. This framework reveals new roles for established *Mtb* virulence determinants at the community scale. The biofilm state confers broad antibiotic tolerance, with PDIM-dependent higher-order organization providing additional protection against isoniazid. By impairing biofilm establishment, EGCG provides proof of principle for chemical perturbation of this community state and highlights biofilm establishment as a potential therapeutic vulnerability in tuberculosis^49,58^.

## ACKNOLEDGEMENTS

We thank the Core Facility Imaging at the Medical University of Vienna for technical assistance and the Center for Microscopy and Image Analysis at the University of Zurich for assistance with transmission electron microscopy. Confocal imaging was performed using equipment maintained by both facilities. We acknowledge the Infectious Diseases Imaging Platform at Heidelberg University, which is funded by the Federal Ministry of Education and Research and the Ministry of Science Baden-Württemberg within the framework of the Excellence Strategy. We further acknowledge the Research Platform Medical Imaging at the Medical University of Vienna for fostering interdisciplinary exchange across imaging disciplines. We thank Baubak Bajoghli of Austrian BioImaging for valuable discussions and continued scientific exchange on imaging-related topics.

## ONLINE METHODS

### Bacterial Strains

All avirulent *Mycobacterium tuberculosis* strains used in this study were derived from the H37Rv background. The PDIM+/ΔRD1 (AE1601) and PDIM−/ΔRD1 (AE1611) strains were clonal PDIM-positive and PDIM-negative isolates, respectively, selected from H37Rv mc² 6230 (ΔRD1 Δ*panCD*)^51^. The PDIM−/ΔRD1 strain additionally carries a ppsC c.2685(C)7→8 frameshift mutation resulting in PDIM deficiency. The PDIM+/RD1+ (AE1201) and PDIM−/RD1+ (AE1219) strains were clonal PDIM-positive and PDIM-negative isolates, respectively, selected from H37Rv mc² 6206 (ΔleuCD ΔpanCD). The PDIM−/RD1+ strain carries a fadD26 nonsense mutation (Trp43*) resulting in PDIM deficiency. Throughout the manuscript, these strains are referred to by PDIM and RD1 status, with RD1+ denoting an intact RD1 locus and ΔRD1 denoting deletion of this locus. Virulent *Mtb* Erdman wild-type and PDIM-deficient 5′Tn::fadD26 strains were used for BSL-3 experiments^70^.

### Culture Conditions

All avirulent strains were routinely cultured using Middlebrook 7H9 base (0.2% glycerol, 0.05% tyloxapol, 10% OADC (oleic acid, bovine serum albumin, dextrose, catalase, sodium chloride)) or Middlebrook 7H10 (0.5% glycerol, 10% OADC) on solid agar base. The cultures were supplemented with 24 µg/ml calcium pantothenate and 50 µg/ml L-leucine as needed; PDIM-positive cultures were additionally supplemented with sodium propionate (0.1 mM in liquid, 1 mM in agar).

Culturing of all BSL-3 mycobacterial strains was performed at 37°C in Middlebrook 7H9 base (Difco) supplemented with 0.5% albumin, 0.2% glucose, 0.085% sodium chloride, 0.5% glycerol, 0.02% tyloxapol in 30 mL inkwell bottles with shaking at 200 rpm.

### Biofilm Induction

To induce biofilm formation in vitro, logarithmic phase bacterial cultures were centrifuged at 3500 ×g for 10 minutes at room temperature. The cell pellet was washed once with biofilm medium (Middlebrook 7H9 base, 0.2% glycerol, 5% OADC) and cell density was adjusted to OD_600_ 1.0 in the same medium. Supplements (calcium pantothenate, L-leucine, and sodium propionate) were added as necessary. Dithiothreitol (DTT) was added to a final concentration of 6 mM, and cultures were mixed thoroughly before being dispensed into the indicated culture vessels. Samples for fluorescence confocal microscopy were cultured in ibidi µ-Slide 8 Well high plates. For crystal violet biomass measurements, 500 µl of OD₆₀₀-adjusted culture was dispensed into each well of a 24-well plate, with three replicate wells prepared per condition. Parallel cultures were incubated with or without 6 mM DTT for 72 h under static, humidified conditions at 37 °C. For mid-IR microscopy, autoclaved Kevley slides were aseptically placed in sterile 100 mm × 15mm petri dishes. 300 µl of culture was dispensed per well in the ibidi slides, and 25 mL in each petri dish. The plates or slides were incubated at 37 °C in static, humidified conditions.

### Biofilm Fixation

Freshly prepared 4% paraformaldehyde (PFA) was used to fix *Mycobacterium tuberculosis* biofilms. The culture media was first carefully removed, then washed twice with DPBS (Corning, 21-031-CV). After DPBS was removed, 4% PFA was dispensed to completely submerge the biofilm and incubated for one hour at room temperature. Following fixation, the fixing solution was removed, and samples were washed twice with sterile water and stored in DPBS at 4°C, protected from light, until staining. Samples for mid-IR imaging were air dried overnight before storing at room temperature, away from light, until ready for measurement. The fixation protocol was validated prior to the removal of samples from the BSL-2 certified laboratory for downstream imaging steps. PFA-treated biofilms were washed twice with sterile water and then scraped off from the wells using wide-bore pipette tips. The entire content was then plated on agar plates (Middlebrook 7H10, 0.5% glycerol, 10% OADC, 24 µg/ml calcium pantothenate and 50 µg/ml L-leucine, 1 mM sodium propionate). The agar plates were incubated at 37 °C for 4 weeks to confirm that no viable bacteria were present.

### Quantification of Biofilm Biomass by Crystal Violet Staining

Following a 72h induction period, supernatant in each well containing *Mtb* biofilm was carefully removed by pipetting. The biofilms were washed twice with sterile DPBS before adding 500 µl of 0.1% crystal violet solution into each well. The wells were incubated with the stain for 30 minutes at room temperature. After 30 minutes, the staining solution was removed and washed thrice with DPBS to remove excess dye. To extract biofilm-bound crystal violet, 500 µl 95% ethanol was added to destain for 30 minutes at room temperature. A 200 µl aliquot of each ethanol extract was transferred to a flat-bottomed 96-well plate, and absorbance was measured at 595 nm. Extracts from DTT-treated samples shown in Extended Data Fig. 1b were diluted 20-fold before measurement, and the reported values were corrected for this dilution. The experiment was performed with three independently grown biological replicates per condition and repeated in an independent experiment with comparable results.

### Enzymatic Treatment of Biofilms

Following a 72h induction period, supernatant in each well containing *Mtb* biofilm was removed by pipetting. The biofilms were washed twice with sterile DPBS. For DNase treatment, the buffer was prepared by diluting the 10X buffer provided in the Turbo DNase kit with sterile water, and DNase was diluted to 0.8 U/mL. Proteinase K was added to DPBS (pH 7.4) to make a final concentration of 0.1 mg/mL. Cellulase and pullulanase were diluted in 0.1 M sodium acetate buffer at pH 5 to 120 mg/mL and 4 NPUN/mL respectively. 500 µl of the enzyme mix was aliquoted into each well containing the washed biofilms. The plates were then incubated for 24 hours at 37 °C. After 24 hours, the supernatant was sampled for the quantification of glucose. The remaining supernatant was removed, and the biofilms were washed twice with sterile DPBS before crystal violet staining.

Enzymatic treatment experiments (Fig. 4b,d) were performed with three independently grown biological replicates per condition and repeated in an independent experiment with comparable results.

### DNS (3,5 –dinitrosalicylic acid) Colorimetric Assay for Reducing Sugar Quantification

The DNS solution was prepared as such: 1.0 g of DNS was added to 50 ml of distilled water and heated to 90°C until dissolved. The solution was allowed to cool, and 20 ml of 0.1 g/ml NaOH was added. 30 g sodium potassium tartrate was added gradually to the solution. Finally, the volume was adjusted to 100 ml using distilled water.

50 µl of a sample (biofilm supernatant or glucose standards) was added to equal volume of the DNS solution and heated at 95°C for 10 minutes. 50 µl of the cooled samples were then added to 50 µl of water, and the absorbance at 540 nm was then measured using a spectrophotometer.

### Formic acid treatment of biofilms

Following a 72 h induction period, supernatant was removed from each well and biofilms were washed twice with sterile DPBS. Biofilms were then exposed to DPBS control or to 88%, 70%, or 50% formic acid for 1.5-2 min at room temperature. The treatment solution was removed, and biofilms were washed twice with DPBS before crystal violet staining and biomass quantification as described above. The biomass experiment shown in Fig. 4e,f comprised three independently grown biological replicates per condition.

For confocal imaging, biofilms treated with 88% formic acid were washed twice with DPBS and neutralized with 1.0 M Tris. Samples were subsequently washed three times with PBS, and the final wash was confirmed to be neutral using pH indicator strips before ThT staining.

### Minimum inhibitory concentration determination

Isoniazid, rifampicin, and (-)-epigallocatechin gallate (EGCG) were solubilized in dimethyl sulfoxide (DMSO). The compounds were serially diluted and 2 μl was spotted onto each well on a 96-well flat bottom plate. 2 μl DMSO was spotted onto solvent-control wells. Mid-log *Mtb* cultures were diluted to OD_600_ 0.005, and 200 μl culture was dispensed into each well. EGCG develops coloration over time, hence bacteria-free controls containing EGCG and media were included for background normalization. Light absorbance at 600 nm was measured in the spectrophotometer after 10 days incubation in a humidified incubator shaking at 100 rpm. The experiment in Extended Data Fig. 7 was independently repeated with three biological replicates yielding comparable results.

### EGCG treatment of biofilms

To assess the effect of EGCG on biofilm formation and established biofilms, (-)-epigallocatechin gallate (EGCG) was added either at the time of biofilm induction together with 6 mM DTT or after 72 h of biofilm maturation. For induction treatment, EGCG was added to a final concentration of 250 µg/ml before dispensing cultures into 24-well plates. For treatment of established biofilms, 72 h biofilms were washed once with sterile DPBS and supplied with fresh biofilm medium containing 250 µg/ml EGCG. Control wells received the corresponding DMSO concentration (1% v/v). Biofilms were incubated for a further 4 days before biomass was quantified by crystal violet staining as described above. The biomass experiment shown in Fig. 4h,i was independently repeated with three biological replicates yielding comparable results.

For confocal imaging, biofilms were exposed to 250 µg/ml EGCG together with DTT for 72 h. The supernatant was subsequently removed, and the retained biofilm material was washed twice with PBS before ThT staining.

### Sample preparation for negative-stain electron microscopy

Biofilm material for negative-stain EM was harvested using a wide-bore pipette tip and transferred to a 1.5 ml tube. Samples were centrifuged at 13,000 × g for 20 min. After removal of the supernatant, the matrix-containing material was collected using a spatula and resuspended in fresh biofilm medium. For analysis of proteinase K-resistant material, biofilms were incubated with 0.1 mg/ml proteinase K for 24 h, as described above. The supernatant was subsequently transferred to a fresh tube without disturbing large detached biofilm fragments and processed for negative-stain EM.

### Bacterial viability assay in antibiotic challenge

*Mtb* biofilms in 24-well plates were prepared as described before, with the exception that the cell density was adjusted to OD_600_ 0.5 and 0.5 ml of culture was dispensed into each well. After 72 hours incubation, the biofilm media containing DTT was removed and the biofilms were washed once with sterile DPBS. Fresh biofilm medium (without DTT) containing various antibiotic or solvent (DMSO) was added to respective wells. For suspension cultures, mid-log *Mtb* cultures were centrifuged at 3,500 ×g for 10 minutes. The cell pellet was then washed once with the fresh medium and adjusted to cell density of OD_600_ 0.5. 5 ml of culture was dispensed into square 30 ml PETG bottles. DMSO, isoniazid, or rifampicin was added into each bottle (n=3 biological replicates per condition). Isoniazid was used at 1.7 or 8.5 µM, and rifampicin was used at 0.4 or 2.0 µM. The final concentration of DMSO in the assay media was limited to 1% to prevent solvent-related toxicity effects. The biofilms were incubated in a humidified standing incubator, while the suspension cultures were kept shaking at 100 rpm. Bacterial viability was determined after 3, 6, and 9 days of incubation. 3 wells of biofilms from each condition were sacrificed at every timepoint: the biofilm was thoroughly scraped from the bottom of the well using a wide-bore pipette tip, and the entire contents transferred into a 1.5 ml snap cap tube. The tubes were centrifuged at 3,500 ×g for 10 minutes, and the supernatant was removed. 0.25 ml of PBS was added to each tube to resuspend the biofilm pellet, and briefly sonicated (10 seconds) in a water bath sonicator. 250 µl of suspension culture was sampled from each bottle and placed in 1.5 ml snap cap tubes. The tubes were centrifuged, resuspended and sonicated in the same workflow. The extracts were then serially diluted and plated on 7H10 agar plates with all necessary supplements. Bacterial viability was determined by enumerating the colony forming units after 21 days and accounting for the corresponding dilution factor.

### Preparation of Staining Reagents

Concanavalin A, Alexa Fluor™ 647 Conjugate stock solution (2 mg/mL) was prepared using 0.1 M sodium bicarbonate buffer (pH 8.5) and stored at-20 °C. Working solution is prepared fresh by diluting the stock solution 1:9 in DPBS containing calcium and magnesium. Working solution of Fluorescent Brightener 28 is prepared fresh by diluting 25% (w/v) Fluorescent Brightener 28 Disodium Salt Solution (Sigma Aldrich) in PBS to 1 mg/mL. The working solution of NucGreen stain was prepared fresh by adding 2 drops of NucGreen™ Dead 488 ReadyProbes™ Reagent to 1 mL of PBS, pH 7.4. A 1 mM Thioflavin T (ThT) stock solution was prepared using 50 mM Tris-HCl (pH 8.0). The solution was passed through a 0.2 µm syringe filter to remove any aggregates and stored in aliquots at-20°C, protected from light. ThT working solution was prepared fresh by diluting the stock solution to 25 µM using 50 mM Tris-HCl.

### Sequential Fluorescent Staining of the Biofilm

All staining and wash steps were performed at room temperature (approx. 20-25°C) and protected from light. Volumes of 200 µL per well were used for staining and washing in ibidi µ-Slide 8 Well high plates, unless otherwise indicated. Imaging commenced approximately 15 minutes after completion of the final staining step.

### Sequential Multi-component Staining

#### NucGreen™ Dead 488 Staining (Nucleic Acids)

The PBS overlying fixed biofilms was removed. Biofilms were washed once with PBS. After aspirating the wash, NucGreen™ Dead 488 working solution was added to each well and incubated for 10 minutes. The staining solution was removed, and wells were washed 3 times with PBS.

#### Fluorescent Brightener 28 Staining (β-Polysaccharides)

Following the final PBS wash from NucGreen staining, PBS was removed. FB28 working solution (1 mg/mL) was added to each well and incubated for 10 minutes. The staining solution was removed, and wells were washed 3 times with DPBS.

#### Concanavalin A, Alexa Fluor™ 647 Staining (α-Polysaccharides)

Following the final DPBS wash from FB28 staining, DPBS was removed. After aspirating the final DPBS wash, ConA-647 working solution (200 µg/mL) was added to each well and incubated for 30 minutes. The staining solution was removed, and wells were washed 3 times with DPBS.

#### Sample Preparation for Imaging

After the final wash, 200 µL of fresh DPBS was left in each well to keep the samples hydrated during microscopy.

#### Thioflavin T staining

For ThT staining, fixed biofilms were stained separately. The PBS overlying the fixed biofilms was removed, and the wells were washed 3 times with 50 mM Tris-HCl buffer. After aspirating the wash, 200 µL of the 25 µM ThT working solution was added to each well and incubated for 30 minutes. The staining solution was subsequently removed, and the wells were washed three times with 50 mM Tris-HCl buffer. After the final wash, 200 µL of fresh 50 mM Tris-HCl buffer was left in each well to keep the samples hydrated during microscopy.

## MICROSCOPY AND IMAGE ACQUISITION

### Transmission Electron Microscopy imaging

Biofilm samples were fixed by immersion in 2.5% Glutaraldehyde in 0.1M cacodylate buffer (pH 7.4). Samples were washed 3 times for 10 min in 0.1 M cacodylate buffer (pH 7.4), followed by post-fixation for 1 h on ice in reduced osmium solution prepared from potassium ferrocyanide, 0.1 M cacodylate buffer, and 2% osmium tetroxide. After washing in distilled water, samples were incubated for 40 min in freshly prepared thiocarbohydrazide solution, washed again, and post-fixed in 2% osmium tetroxide for 90 min at room temperature. Samples were then washed and stained overnight in 1% aqueous uranyl acetate at 4 °C. On the following day, samples were washed in distilled water, including three 10-min washes in pre-warmed water at 50 °C, and incubated for 2 h at 50 °C in lead aspartate solution adjusted to pH 5.0. After washing, samples were dehydrated through a graded ethanol series (70%, 80%, 96%, and 100%; 20 min each), followed by two additional 15-min washes in absolute ethanol and two 15-min washes in propylene oxide. Samples were infiltrated overnight in a 1:1 mixture of Epon/Araldite and propylene oxide, transferred to 100% Epon/Araldite for 50 min, flat-embedded in fresh Epon/Araldite in Aclar foil, and polymerized at 60 °C for 28 h. Thin sections (70nm) were imaged in a Talos 120 transmission electron microscope at 120 kV acceleration voltage equipped with a bottom mounted Ceta camera using the Maps software for automatic image acquisition (Thermo Fischer Scientific, Eindhoven, The Netherlands).

For the sectioned TEM analysis shown in Extended Data Fig. 3, one independently grown biofilm sample was analysed per genotype, with at least three spatially distinct regions examined within each sample.

For negative stain EM analysis of biofilm matrix or proteinase K-treated biofilm material, 4µl of the sample were applied to glow-discharged, carbon-coated grids for 40 s, washed twice with water, once with 1% phosphotungstic acid and subsequently stained with 1% phosphotungstic acid for 40 s. The grids were examined using a Morgagni transmission electron microscope (Thermo Fischer Scientific, Eindhoven, The Netherlands) operated at 80 kV.

Negative-stain EM was performed on two independently prepared biofilm samples generated on separate days, with at least three spatially distinct regions examined per sample.

### Fluorescence imaging

Confocal microscopy was performed using a Leica TCS SP8 FALCON point-scanning confocal microscope system, built on a Leica DMI8 inverted microscope stand (Leica Microsystems). The system was equipped with a 405 nm solid-state laser, a pulsed White Light Laser (WLL; 470–670 nm tunable range), and Hybrid Detectors (HyD). Image acquisition was controlled using Leica Application Suite X (LAS X) software.

Imaging was conducted using a 63x immersion microscope objective (NA=1.3). Images were acquired with a resolution of 2048 x 2048 pixels (184.52 x 184.52 µm field of view; 90.14 x 90.14 nm pixel size) and 3x line averaging, using a scan speed of 600 Hz and a pinhole setting of 0.85 AU. Three-dimensional (3D) reconstructions were generated from Z-stacks acquired with a 1 µm step size through the depth of the biofilm.

For multi-component imaging, two separate acquisitions were performed in frame sequential mode to prevent spectral bleed-through. The first acquisition captured the polysaccharide signals, and the second captured the nucleic acid signal from the same field of view. The settings were as follows:

- **Calcofluor White (FB28):** Excitation: 405 nm. Emission detection: 430–500 nm.
- **Concanavalin A, Alexa Fluor™ 647 (ConA-647):** Excitation: 653 nm (from WLL). Emission detection: 658–775 nm.
- **NucGreen™ Dead 488 (SYTOX™ Green):** Excitation: 504 nm (from WLL). Emission detection: 510–630 nm.

For separate ThT imaging experiments, the following settings were used:

- **Thioflavin T (ThT), bound:** Excitation: 405 nm. Emission detection: 480–530 nm.
- **Thioflavin T (ThT), free control:** Excitation: 405 nm. Emission detection: 415–460 nm.

Laser power for each channel was adjusted to avoid detector saturation. Typical laser power settings were: FB28 at 8%, NucGreen at 20%, ConA-647 at 1.7%, and ThT at 8%. HyD detector gain was generally set to 80%. All imaging was performed in a dark room, commencing approximately 15 minutes after the final staining wash.

Confocal microscopy of PFA-inactivated, BSL-3 *Mtb* biofilms was performed on an inverted Leica SP8 point laser scanning confocal microscope using an HC PL APO CS2 63x/1.4 oil immersion objective and the LasX microscope-control software. Images were acquired with 405 nm laser excitation. Laser power and detector gain was generally adjusted to avoid signal saturation. Bound and free ThT detection settings were setup as mentioned previously. Images are representative of two independently grown biological replicates per background, with at least three spatially distinct regions imaged per biofilm.

For the multicomponent fluorescence imaging shown in Fig. 1 and Extended Data Fig. 2, three biological replicates per genotype were analysed in each of two independent experiments, with at least three spatially distinct regions imaged per biofilm and comparable genotype-dependent architectures observed across experiments. For the genotype-comparison ThT imaging shown in Fig. 4g, three biological replicates per background were examined in each of two independent experiments, with at least three spatially distinct regions imaged per biofilm. For the proteinase K-detached material shown in Extended Data Fig. 4, three biological replicates were analysed, with at least three spatially distinct fields imaged per biological replicate and comparable cord-like structures observed across samples.

### QCL based Mid-Infrared (Mid-IR) Reflection Microspectroscopy

Biofilms representing the PDIM+/RD1+, PDIM+/ΔRD1, PDIM−/RD1+ and PDIM−/ΔRD1 genotype backgrounds were cultured and fixed as described above and prepared for mid-IR imaging. Briefly, after fixation in 4% paraformaldehyde and rinsing with sterile distilled water, biofilm-coated mid-IR-compatible MirrIR low-e microscope slides (Kevley Technologies) were aseptically air-dried and stored at room temperature until analysis^68^.

Hyperspectral mid-IR imaging was performed using a Quantum Cascade Laser (QCL)-based mid-IR microscope (Spero® QT 340, DRS Daylight Solutions) operating in reflection mode. The system is equipped with a tunable QCL source covering the fingerprint region from approximately 1800 cm⁻¹ to 950 cm⁻¹ and a 480×480 focal plane array (FPA) detector, enabling a spatial resolution of 12 µm at a wavelength of 5.5 µm as well as rapid data acquisition (∼1 minute per hyperspectral cube). Reflection mode was chosen for its suitability with fixed biofilm samples on the reflective Kevley slides.

Hyperspectral data cubes were acquired using the SperoChemVision™ software (Daylight Solutions). Data processing was conducted using a custom Python-based pipeline. To mitigate baseline variability arising from sample thickness heterogeneity and optical scattering effects, spectra at each pixel were z-score normalized: each spectrum’s mean intensity was subtracted, and the result was divided by its standard deviation across the 950-1800 cm⁻¹ range. No further baseline correction algorithms were applied post-z-score normalization.

For the mid-IR analyses shown in Figs. 2 and 3, one independently grown biofilm was analysed per genetic background, with at least three spatially distinct regions measured per biofilm.

## DATA PROCESSING

Color overlay images were generated using Fiji (ImageJ). For merged images presented in Figure 1d, h, l and p, a threshold was applied to the lower intensity range of the green channel (nucleic acid signal) to enhance the visualization of brighter bacilli against any diffuse background signal.

### Generation of Thresholded Specific Wavenumber Abundance Maps (TSWAMs)

To visualize the spatial distribution of specific biomolecular signatures, TSWAMs were generated from the z-score-normalized hyperspectral data. Wavenumbers were selected from prominent local spectral maxima within the corresponding biomolecular regions for each dataset. Consequently, closely related spectral features could be represented by slightly different wavenumbers between genetic backgrounds, for example 1450 or 1460 cm⁻¹ within the lipid/mixed-signal region.

For each selected wavenumber, intensity values across all pixels in the two-dimensional image were linearly scaled to a range of 0-1. A binary mask was then generated using an intensity threshold of >0.55. Within this mask, the scaled intensity values were re-normalized to 0-1, such that the retained intensity range (for example 0.55-1) spanned the full display range. This second scaling enhanced the dynamic range of the retained signal for visualization without altering which pixels were included in the mask.

TSWAMs therefore represent regions of higher relative signal at the selected wavenumber and were used for qualitative assessment of biochemical distribution and compartmentalization. The approximate field of view was 1.4 × 1.4 mm.

### Principal Component Analysis (PCA) of Mid-IR Hyperspectral Data

Prior to clustering, Principal Component Analysis (PCA) was applied to the z-score normalized mid-IR hyperspectral data cubes (950-1800 cm⁻¹, one spectrum per pixel) for each *Mtb* strain to reduce dimensionality and noise. PCA was performed using sklearn.decomposition.PCA. The number of principal components (PCs) retained for subsequent clustering was determined by the Kaiser criterion, i.e., selecting PCs with eigenvalues greater than 1. The retained PC scores were then standardized using StandardScaler from scikit-learn before K-means clustering.

### K-means Clustering and Cluster Evaluation

K-means clustering was performed on the retained PCs for each *Mtb* strain dataset using the KMeans function from sklearn.cluster. The algorithm was run with 10 different initializations (n_init=10) and a fixed random state (random_state=42) for reproducibility, selecting the run with the lowest inertia. To determine the optimal number of clusters (k) for each strain, a range of k values from 2 to 15 was evaluated using several common clustering metrics:

1. **Inertia (Elbow Method):** The sum of squared distances of samples to their closest cluster center was plotted against k to identify an “elbow” point, suggesting a suitable k.
2. **Davies-Bouldin Score:** This metric measures the average similarity ratio of each cluster with its most similar cluster. Lower values indicate better clustering, with a minimum score of 0.
3. **Calinski-Harabasz Index (Variance Ratio Criterion):** This index is the ratio of the sum of between-cluster dispersion to within-cluster dispersion. Higher values generally indicate better-defined clusters.

The final number of clusters for each genotype background (PDIM+/RD1+, k = 4; PDIM+/ΔRD1, k = 3; PDIM−/RD1+, k = 3; PDIM−/ΔRD1, k = 3) was selected based on consensus across these metrics.

### Generation of Cluster Assignment Maps and Average Spectra

Cluster assignment maps were generated by assigning each pixel in the original spatial dimensions to the cluster determined by K-means analysis of its corresponding PCA-reduced spectrum. Average cluster spectra were then calculated for each identified cluster by averaging the original z-score normalized mid-IR spectra (950-1800 cm⁻¹) from all pixels assigned to that cluster.

### CLSM fluorescence segmentation and quantification

Single-channel 8-bit CLSM z-stacks of untreated, EGCG-treated and formic acid-treated biofilms were analysed. Five biological replicates were imaged in the primary experiment, with one additional independently grown biological replicate imaged on a separate day; at least three spatially distinct regions were imaged per biofilm.

Analyses included 15 optical planes, corresponding to approximately 0–14 µm from the first acquired plane, without alignment to peak fluorescence. All conditions were imaged using identical acquisition settings, and images used for visual comparison were displayed using the same intensity range.

Segmentation was performed separately for each optical plane. Images were smoothed and normalized using their robust intensity range to account for changes in signal intensity across depth. Otsu thresholding was used to identify confidently fluorescent seed regions. Masks were subsequently expanded through connected pixels with normalized intensities of at least 20% of the plane-specific Otsu threshold, thereby incorporating dimmer fluorescence associated with the segmented structures while excluding spatially disconnected signal. Small isolated regions were removed, followed by binary closing and dilation to consolidate adjoining structures. Glass-associated fluorescence in the first two planes was reduced by retaining locally structured signal or signal spatially supported by planes 3 and 4. Segmentation quality was assessed using binary masks, overlays with the original images and residual images showing fluorescence outside the masks.

Fluorescence was quantified from the original unfiltered images without background subtraction. For each optical plane, ThT-positive area fraction was calculated as the number of masked pixels divided by the total number of pixels in that plane. Mean fluorescence within the mask was calculated from the raw intensities of masked pixels. Coverage-weighted fluorescence, reported as area-normalized integrated fluorescence, was calculated as the sum of raw intensities within the mask divided by the total number of pixels in the plane and therefore incorporates both fluorescent coverage and fluorescence intensity.

For stack-level summaries, masked-pixel counts, raw intensity sums and total pixel counts were pooled across all 15 optical planes. ThT-positive volume fraction was calculated as the number of segmented voxels divided by the total number of voxels within the analysed stack. Mean fluorescence within the segmented volume was calculated from the raw intensities of segmented voxels, and volume-normalized integrated fluorescence was calculated as the summed raw fluorescence within the segmented volume divided by the total number of voxels. Stack-level values were expressed relative to the untreated arithmetic mean for visualization.

## STATISTICAL ANALYSIS

For antibiotic-tolerance experiments, CFU/ml values were log10-transformed, and changes from baseline were calculated using the corresponding strain-and growth-state-specific reference at time zero. Three biological replicates were analysed per condition. Biofilm measurements at each time point were obtained from independently grown biological replicates because CFU determination was destructive, whereas the three suspension-culture replicates were sampled serially across time.

Biofilm and suspension conditions were compared separately at each compound concentration and sampling time using ordinary two-way ANOVA with strain and growth state (biofilm or suspension) as fixed factors. Biofilm and suspension means were compared within each strain using simple-effect comparisons with Šídák adjustment. The four biofilm-suspension comparisons across the four strains at each concentration and timepoint constituted one multiple-comparison family.

Differences among strains in the biofilm state over time were analysed separately for each compound concentration using ordinary two-way ANOVA with strain and timepoint as fixed factors. At each timepoint, all pairwise comparisons among the four strains were performed using Tukey’s multiple-comparisons test, with the six strain comparisons constituting one multiple-comparison family.

For depth-resolved ThT fluorescence analysis, each complete 15-plane z-stack was treated as one independent biological observation (n=6 per condition). EGCG-and formic acid-treated stacks were compared separately with untreated stacks using exact two-sided whole-profile permutation tests. Treatment labels were permuted only between complete z-stacks, thereby preserving the correlation between optical planes within each biological replicate. For each comparison, all 924 possible allocations of the 12 stacks into two groups of six were evaluated.

At each optical plane, the difference between treatment and untreated means was divided by its estimated standard error to obtain a studentized difference. The global test statistic was calculated as the sum of the squared studentized differences across all 15 optical planes, providing a single test of the complete depth profile without treating individual optical planes as independent observations. Area-normalized integrated fluorescence was designated the primary endpoint, with Holm correction applied across its two treatment comparisons. ThT-positive area fraction and mean fluorescence within segmented regions were secondary endpoints, with Holm correction applied across their four treatment comparisons.

Because the exact permutation test evaluates a finite number of label allocations, attainable P values are discrete. For the two-sided tests used here, the observed allocation and its complete label reversal yield the same test statistic, giving a minimum attainable unadjusted P value of 2/924 (P = 0.00216). Following Holm correction, the corresponding minimum adjusted P values were 0.0043 for the two primary-endpoint comparisons and 0.0087 for the four secondary-endpoint comparisons.

Where the corresponding whole-profile test was significant, differences at individual optical planes were localized using a permutation-based maxT procedure. For each permutation, studentized treatment differences were calculated at all 15 depths and the largest absolute value was retained. The observed studentized difference at each depth was compared with this distribution of permutation maxima to obtain a maxT-adjusted P value, thereby controlling the family-wise error rate across the 15 depth-wise comparisons. Depths with maxT-adjusted P < 0.05 are indicated in Extended Data Fig. 6.

For stack-level fluorescence summaries, each biological replicate was represented by a single value pooled across the analysed 15-plane volume. Prespecified comparisons of EGCG-treated and formic acid-treated biofilms with untreated biofilms were performed using two-sided Welch’s t-tests, allowing for unequal variances between groups. Statistical testing was performed on the unnormalized stack-level values, whereas normalization to the untreated arithmetic mean was used only for visualization in Extended Data Fig. 6d. P values were adjusted using the Holm procedure across the six planned comparisons comprising two treatments and three fluorescence measurements.

All statistical tests were two-sided, and multiplicity-adjusted P < 0.05 was considered statistically significant. No statistical method was used to predetermine sample size. No biological replicates were excluded from the reported analyses.

## SOFTWARE

Mid-IR data processing and analysis used Python 3.12.3 with NumPy 1.26.4, SciPy 1.13.0, Matplotlib 3.8.4, PyQt5 5.15.10 and scikit-learn 1.6.1. CLSM triple-stain and ThT image processing and visualization, as well as fibril measurements, were performed using Fiji 1.54p. Depth-resolved ThT fluorescence segmentation, quantification and statistical analysis used Python 3.14.4 with NumPy 2.5.1, SciPy 1.16.1, Pillow 12.3.0, tifffile 2026.7.14 and readlif 0.6.6; figures generated during this analysis used Matplotlib 3.11.1. Statistical analyses of antibiotic-tolerance experiments were performed using GraphPad Prism version 11.0.2. Confocal image acquisition was performed using Leica Application Suite X (LAS X; Leica Microsystems).

## DATA AVAILABILITY

Data supporting the findings of this study are available to editors and reviewers upon request during peer review. Upon acceptance, the minimum dataset necessary to interpret, verify and extend the reported findings will be deposited in a suitable public repository. Repository identifiers and access information will be added before publication.

## CODE AVAILABILITY

No previously unreported custom algorithms central to the conclusions of this study were developed. Mid-IR and depth-resolved fluorescence analyses used established algorithms implemented through standard Python libraries and paper-specific research software. Code required for evaluation of the reported analyses can be made available to editors and reviewers upon request during peer review. Requests for paper-specific research code thereafter will be considered by the corresponding authors upon reasonable request, subject to applicable intellectual-property and software-licensing restrictions.

---

**Extended Data Figure 1:**
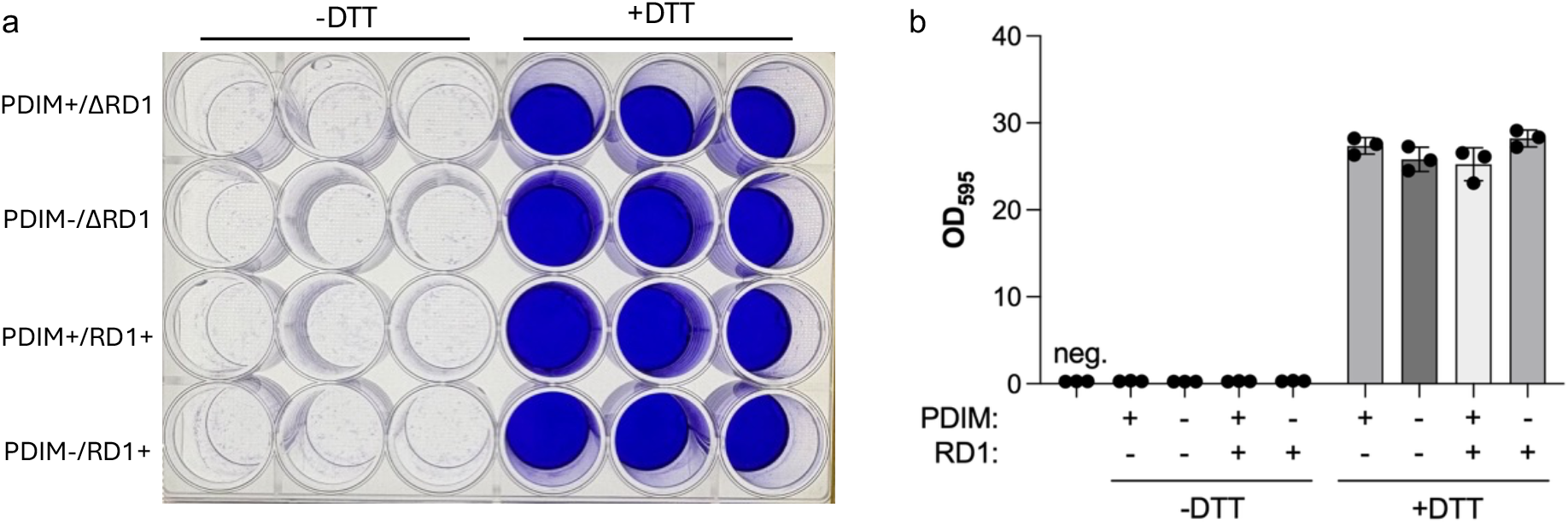
DTT induces submerged *Mtb* biofilm formation independently of PDIM and RD1. a, Representative top view of a crystal violet-stained 24-well plate containing the H37Rv-derived auxotrophic PDIM+/ΔRD1, PDIM−/ΔRD1, PDIM+/RD1+ and PDIM−/RD1+ backgrounds after 72 h of incubation. Columns 1-3 were cultured without DTT, whereas columns 4-6 received 6 mM DTT. **b,** Quantification of retained biofilm biomass by crystal violet solubilization and absorbance measurement at 595 nm. Values from DTT-treated samples were corrected for the 20-fold dilution used for measurement. Data are presented as mean ± s.d. of three biological replicates.

**Extended Data Figure 2:**
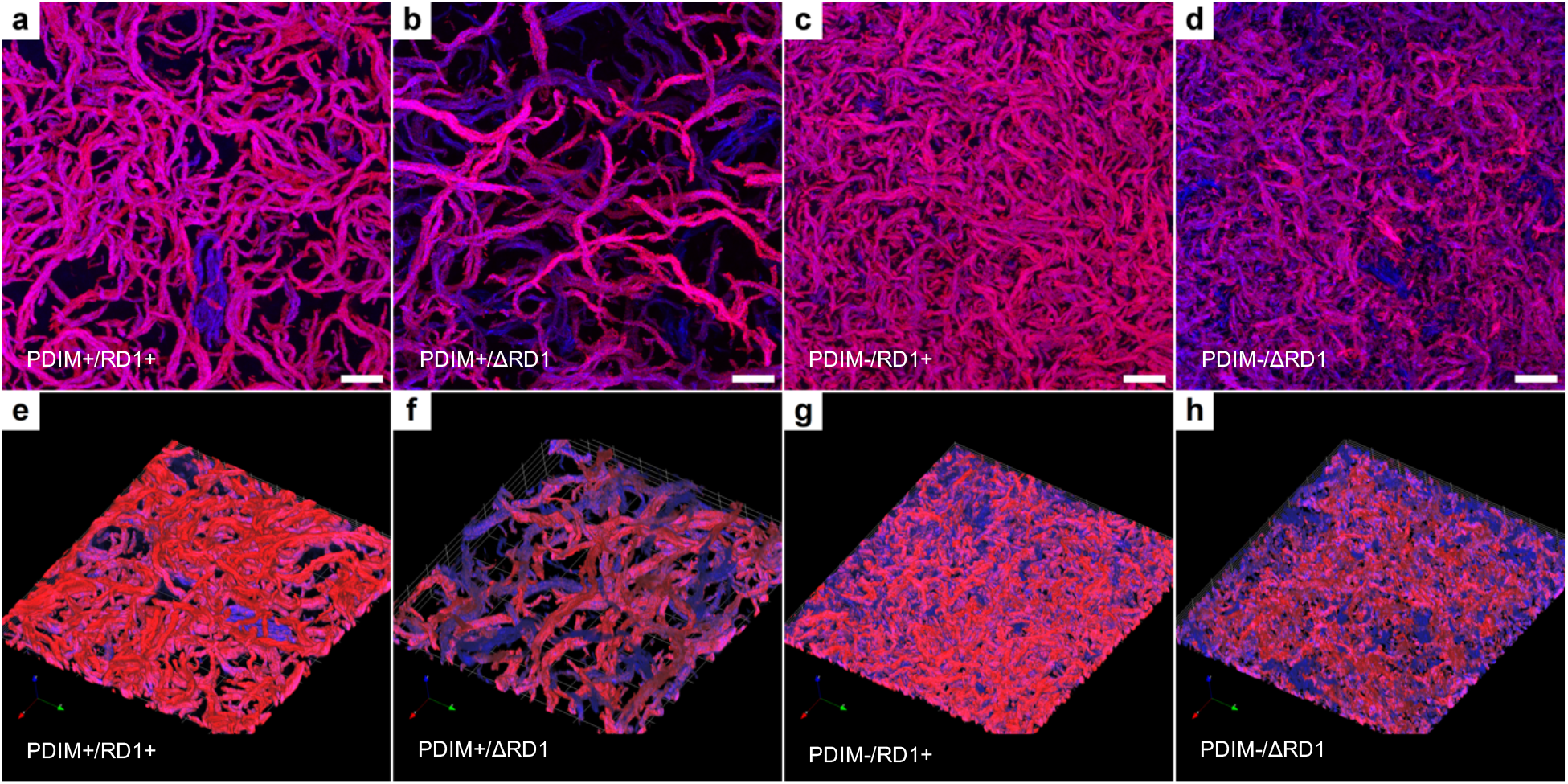
PDIM promotes the higher-order organization of cord-like biofilm structures. The α-polysaccharide (red) and β-polysaccharide (blue) channels from the biofilms shown in Fig. 1 were combined to visualize matrix organization. **a–d,** En face maximum-intensity projections. **e–h,** Corresponding three-dimensional volume renderings. Images show PDIM+/RD1+ (a,e), PDIM+/ΔRD1 (b,f), PDIM−/RD1+ (c,g) and PDIM−/ΔRD1 (d,h) backgrounds. Scale bars, 20 µm.

**Extended Data Figure 3:**
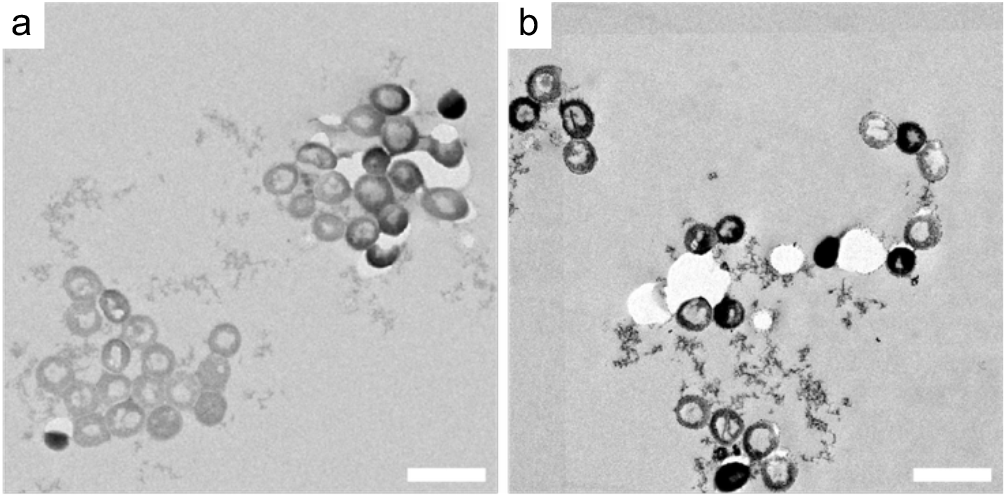
Transmission electron microscopy confirms that cord-like biofilm structures contain aligned bacilli. Representative transmission electron micrographs of sections through formaldehyde-fixed biofilms. **a,** PDIM+/RD1+ biofilm showing closely associated and aligned bacilli. **b,** PDIM−/RD1+ biofilm showing more sparsely distributed bacilli with disordered organization. Bacilli measured approximately 450 nm in diameter. Scale bars, 1 µm.

**Extended Data Figure 4:**
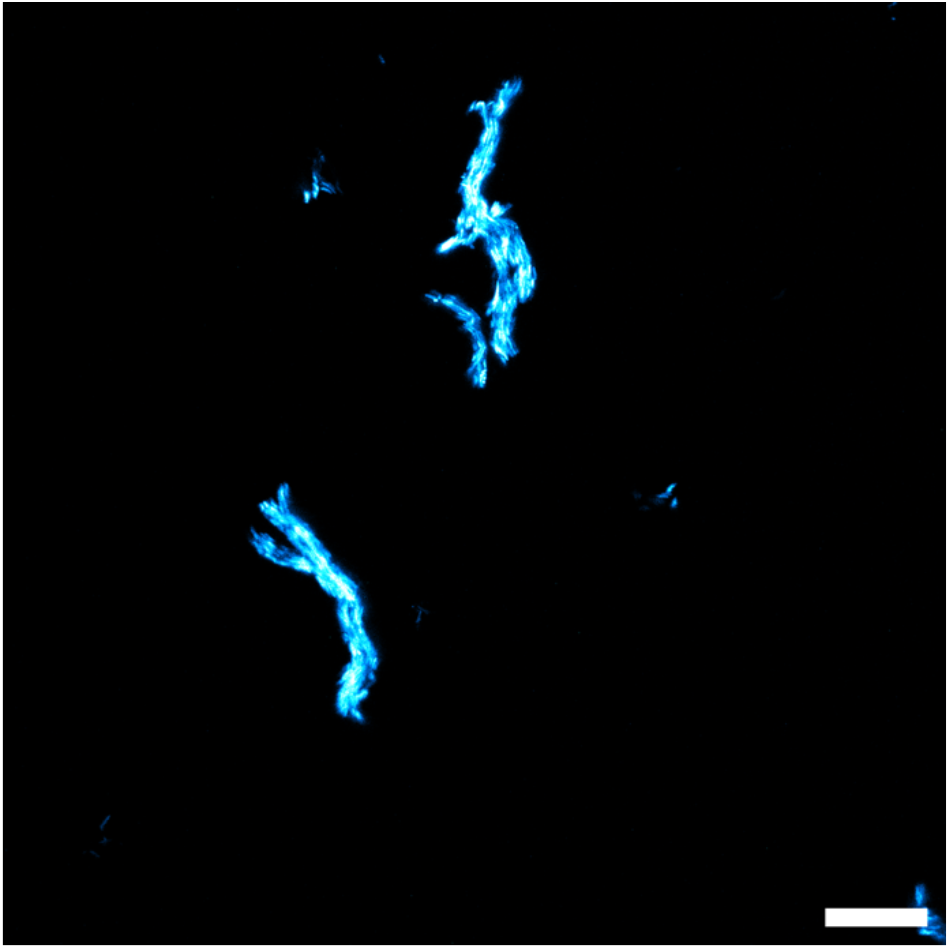
Individual cord structures persist following proteinase K-mediated biofilm detachment. PDIM+/RD1+ biofilms were treated with 0.1 mg/ml proteinase K for 24 h. Detached biofilm material was stained with thioflavin T (ThT) and visualized by confocal microscopy. Scale bar, 20 µm.

**Extended Data Figure 5:**
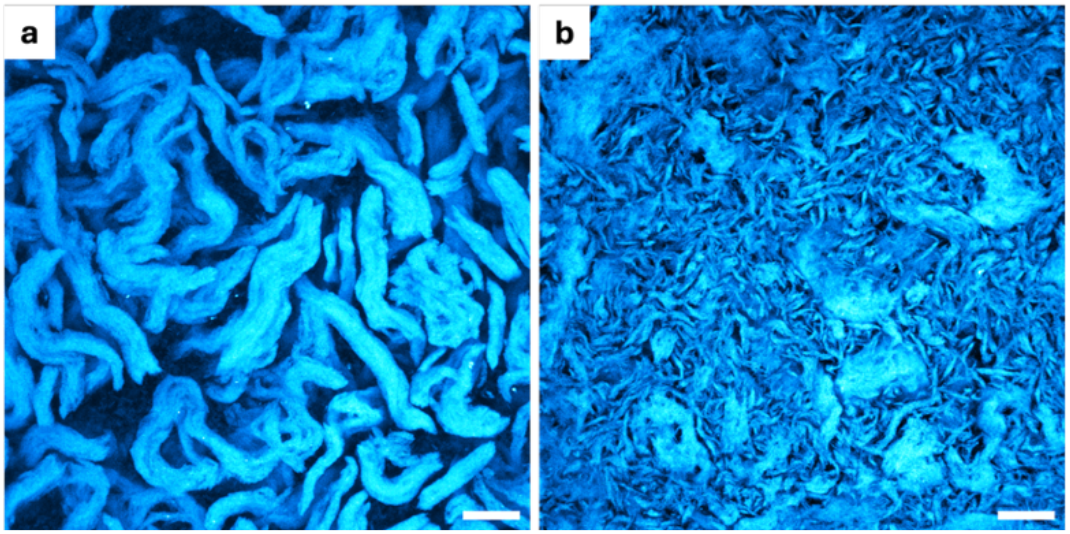
Virulent *Mtb* Erdman biofilms exhibit PDIM-dependent cord organization and ThT-positive matrix material. Representative confocal microscopy images of ThT-stained biofilms formed under DTT-induced thiol reductive stress. **a,** Wild-type Erdman formed thick, serpentine cords associated with ThT-positive matrix material. **b,** The PDIM-deficient Erdman 5′Tn::fadD26 mutant formed shorter, disorganized aggregates while retaining ThT-positive matrix material. Scale bars, 20 µm.

**Extended Data Fig. 6:**
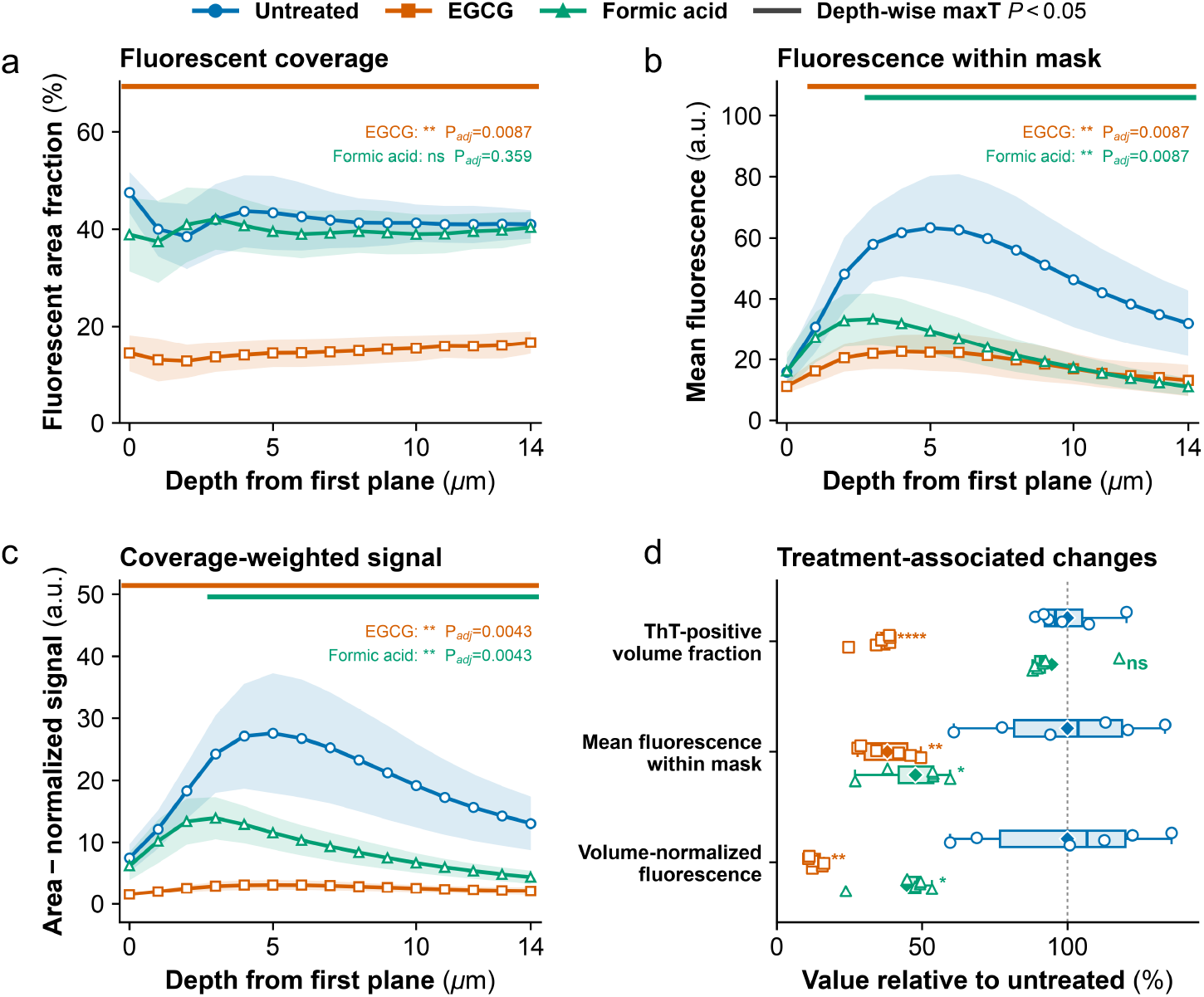
Depth-resolved ThT fluorescence following EGCG or formic acid treatment. Depth profiles of ThT-positive area fraction (**a**), mean raw fluorescence within segmented regions (**b**) and coverage-weighted fluorescence (**c**). Lines show means and shaded regions indicate s.d. across six independent biological replicates. P values are Holm-adjusted values from exact two-sided whole-profile permutation tests; coloured bars indicate depths with maxT-adjusted P < 0.05. **d,** Stack-level ThT-positive volume fraction, mean fluorescence within the segmented volume and volume-normalized fluorescence, expressed relative to untreated. Boxes show the interquartile range with median centre lines; whiskers extend to 1.5 × the interquartile range, open symbols indicate individual biological replicates and filled diamonds indicate means. Significance in d was determined by two-sided Welch’s t-tests on unnormalized values with Holm correction across the six planned comparisons. n = 6 independent biological replicates per condition. ns, P ≥ 0.05; *P < 0.05; **P < 0.01; ***P < 0.001; ****P < 0.0001.

**Extended Data Figure 7:**
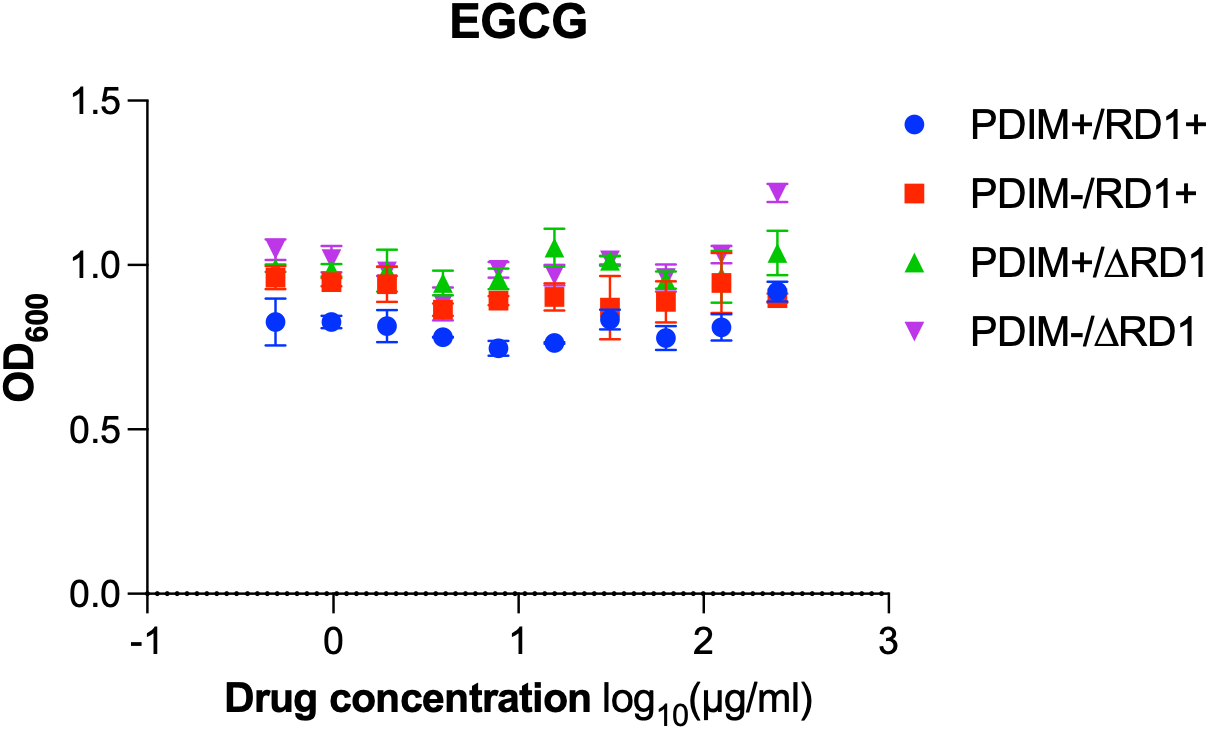
EGCG does not inhibit planktonic *Mtb* growth. PDIM+/RD1+ (blue circles), PDIM−/RD1+ (red squares), PDIM+/ΔRD1 (green triangles) and PDIM−/ΔRD1 (purple triangles) cultures were exposed to the indicated concentrations of epigallocatechin gallate (EGCG) for 10 d. Culture density was subsequently measured by absorbance at 600 nm. Data are presented as mean ± s.d. of three biological replicates per genetic background.

**Extended Data Figure 8:**
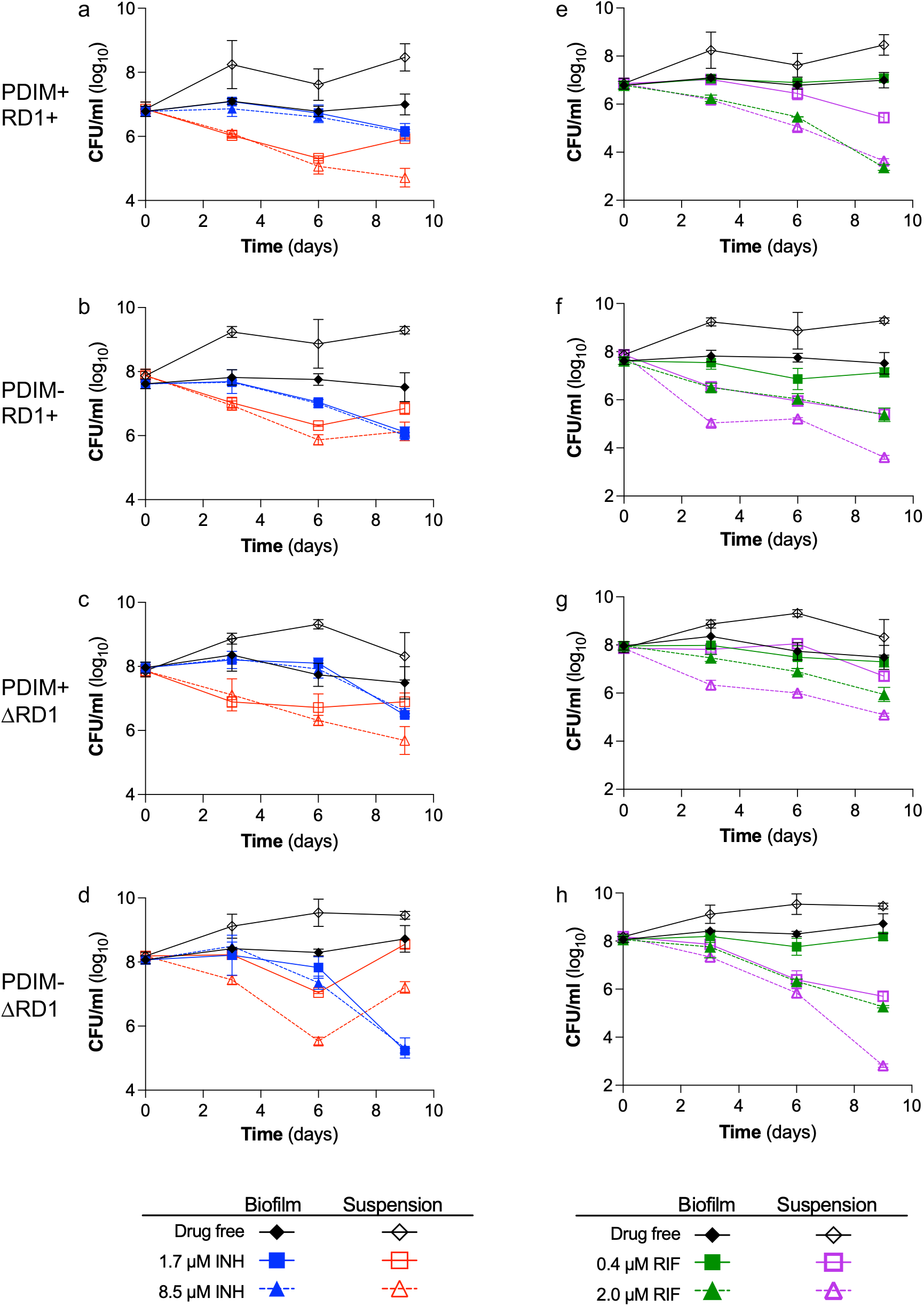
The biofilm state increases *Mtb* tolerance to isoniazid and rifampicin across four genetic backgrounds. Planktonic cultures in 7H9 broth and 3-day-old submerged biofilms were exposed to antibiotics, and viable bacteria were quantified by CFU enumeration at the start of treatment (day 0) and on days 3, 6 and 9. **a–d,** PDIM+/RD1+ (a), PDIM−/RD1+ (b), PDIM+/ΔRD1 (c) and PDIM−/ΔRD1 (d) cultures treated with 1.7 µM (squares, solid lines) or 8.5 µM (triangles, dashed lines) isoniazid. **e–h,** The corresponding backgrounds treated with 0.4 µM (squares, solid lines) or 2.0 µM (triangles, dashed lines) rifampicin. Filled blue or green symbols and lines denote biofilm-associated bacteria, whereas open red or purple symbols and lines denote planktonic cultures. Drug-free controls are shown as black diamonds with solid lines. Data are presented as mean ± s.d. of three biological replicates.

**Extended Data Figure 9:**
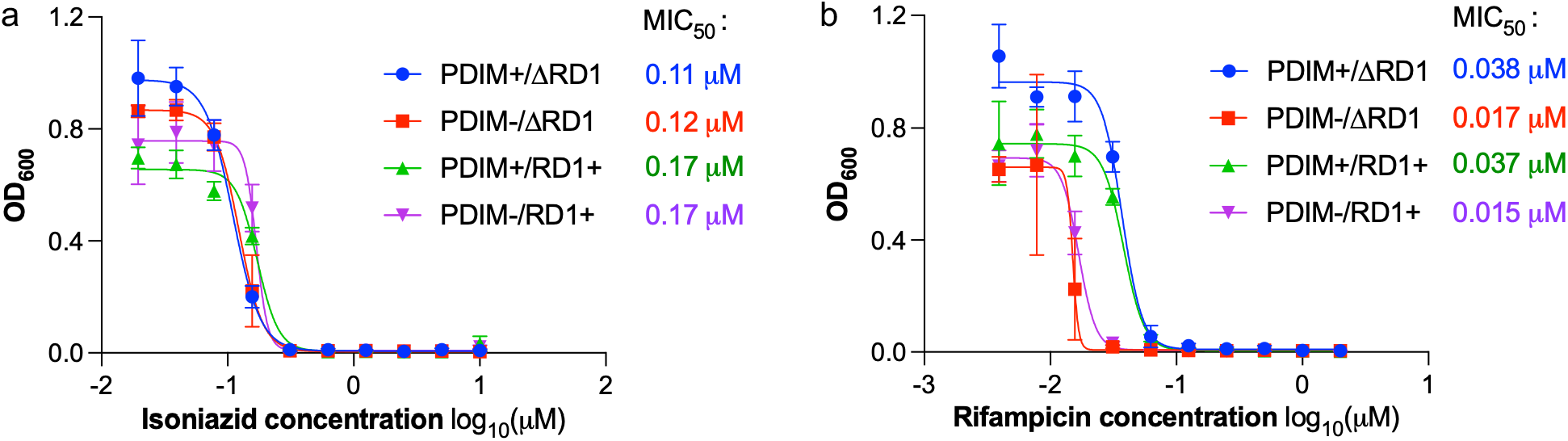
Isoniazid and rifampicin susceptibility across avirulent *Mtb* genetic backgrounds. PDIM+/ΔRD1 (blue circles), PDIM−/ΔRD1 (red squares), PDIM+/RD1+ (green triangles) and PDIM−/RD1+ (purple triangles) cultures were exposed to the indicated concentrations of isoniazid (a) or rifampicin (b) for 10 d. Growth was measured by absorbance at 600 nm. MIC₅₀ denotes the antibiotic concentration required to inhibit growth by 50%. Data are presented as mean ± s.d. of three biological replicates per genetic background.

